# When can neuronal activity-dependent homeostatic plasticity maintain circuit-level properties?

**DOI:** 10.64898/2026.02.07.704433

**Authors:** Lindsay J. Stolting, Randall D. Beer

## Abstract

Neural circuits are remarkably robust to perturbations that threaten their function. Activity-dependent homeostatic plasticity (ADHP) is a stabilizing mechanism that supports robustness by tuning neuronal ion conductances to combat chronic over- or under-activity. Its restorative capacity has been demonstrated in the pyloric circuit of the crustacean stomatogastric ganglion, whose neurons must burst in a specific order to coordinate digestive muscles. After disruption by physical and pharmacological manipulations, this circuit reliably recovers not only the activity levels of constituent neurons, but also the proper burst order. But how could ADHP, operating only on local information about each neuron’s average activity, maintain higher-order circuit properties? We explored this question in a computational model of the pyloric pattern generator. We first optimized a set of pyloric-like networks, then optimized ADHP mechanisms for each network to restore its pyloric character after parametric perturbations. This was possible for some networks and impossible for others, so we aimed to explain this disparity. We found that successful homeostatic regulators target average neural activity levels which happen to occur only among pyloric circuits and not among non-pyloric ones, within the set of reachable circuit configurations. Therefore, in subsets of parameter space where such dissociation is possible, activity carries indirect information about burst order, which ADHP can exploit to maintain pyloricness. Other subsets, whose pyloric averages are inseparable from non-pyloric ones, cannot be perfectly regulated. This separability property may explain differences in recovery capacity across perturbations and across individuals.

## 1 Introduction

Neural circuits are subject to constant changes in their properties and organization as organisms learn, develop, and contend with their environment. Despite these changes, circuits robustly orchestrate behavior throughout an animal’s life-time. Homeostatic neural activity regulation, or homeostatic plasticity, is one important contributor to this robustness (Davis, 2006; Davis & Bezprozvanny, 2001; Turrigiano & Nelson, 2004). Activity-dependent homeostatic plasticity, or ADHP, refers to a family of cellular mechanisms by which neurons sense their own activity levels and regulate intrinsic and synaptic conductances to avoid chronic hyper-or hypo-activity (Davis & Goodman, 1998; Desai, 2003; Northcutt & Schulz, 2019; Pozo & Goda, 2010; Turrigiano, 2012; Wen & Turrigiano, 2024). ADHP has been demonstrated in various experiments where neurons respond to prolonged periods of artificial activity stimulation or suppression with compensatory changes in membrane and synaptic properties. It is often difficult, however, to trace the behavioral consequences of these changes. Therefore, it remains largely unclear whether and how the seemingly simple maintenance of individual neural activity levels could stabilize the intricate neural circuit functions underlying behavior.

For example, ADHP has been observed in cultured rat neurons, which show up- or down-regulation of excitability after activity deprivation or chronic stimulation, respectively (Desai et al., 1999; O’Leary et al., 2010; Turrigiano et al., 1998; Tyssowski et al., 2019). Networks of stem cell derived human neurons show similar responses and showcase mechanistic diversity. Homeostatic responses are correlated in various preparations to extracellular matrix changes, pre-synaptic vesicle localization, post-synaptic receptor distribution, and even astrocyte interaction (Cordella et al., 2022; Yuan et al., 2023). But, while they provide valuable mechanistic data, cultured neurons are divorced from a bodily context and cannot speak to ADHP’s maintenance of coordinated behavior. Conversely, ADHP’s effects have been measured in *in vivo* cortex. In preparations of monocular visual deprivation, the synaptic drive to the input-deprived hemisphere initially decreases, and then steadily increases to compensate for the perturbation over several days (Hengen et al., 2013; Keck et al., 2013; Lambo & Turrigiano, 2013; Maffei & Turrigiano, 2008). It is hypothesized that the purpose of such tuning in sensory cortex is to ensure neural responses are maximally sensitive to incoming stimuli (Aizenman et al., 2003; Stemmler & Koch, 1999; Sullivan & de Sa, 2006; Wen & Turrigiano, 2024), but how do we know that this is sufficient to preserve all the critical functions of the visual cortex? Does the observed compensation affect visual processing operations, or communication with upstream and downstream systems? Or might it have been the case that average activity was restored without these more nuanced properties? In order to investigate homeostatic maintenance of structured behavior, then, it would be best to consider ADHP acting on embodied circuits with well-defined behavioral roles.

One outstanding example, in which ADHP has alreaady been productively studied, is the pyloric subcircuit of the crustacean stomatogastric ganglion. The pyloric circuit is a central pattern generator (CPG) which controls muscles at the passage from the animal’s stomach to the hindgut, directing rhythmic digestive motion (Rezer & Moulins, 1983). It includes an intrinsically bursting kernel, composed of an anterior burster (AB) neuron electrically coupled to two pyloric dilator (PD) neurons. This kernel inhibits two conditionally bursting follower elements: a group of pyloric (PY) neurons and a single lateral pyloric (LP) neuron (Harris-Warrick, 1992). Motor patterns vary within and between individuals, but several dynamical features are remarkably well-conserved (Bucher et al., 2005; Hooper, 1997; Soofi et al., 2012, 2014). Most saliently, circuit elements must burst in a specific order to enact a functional motor rhythm. The conserved repeating sequence is LP—PY—PD/AB, with the pacemaker kernel of PD/AB cells synchronized, and the end of the LP burst slightly overlapping the beginning of the PY burst (Prinz et al., 2004).

When isolated from their synaptic inputs and placed in saline, pyloric neurons initially fall silent and respond to depolarization with tonic spikes, but after three days in culture they respond to the same stimulation with normal-looking burst sequences (Turrigiano et al., 1994, 1995). Likewise, when local connections are preserved but input to the ganglion is removed (either pharmacologically or by severing afferents) the rhythm stops for a time but normal activity is restored over several days (Golowasch, Casey, et al., 1999; Temporal et al., 2012, 2014; Thoby-Brisson & Simmers, 1998; Thoby-Brisson & Simmers, 2000; Zhang & Golowasch, 2011; Zhang et al., 2009). The process of recovery is not uniform and may include multiple intervals where the pyloric rhythm returns and disappears, before eventually stabilizing. These epochs of rhythmic activity are called bouts (Luther et al., 2003).

The ADHP mechanisms at work in this preparation appear to track calcium influx as an indicator of elevated or depressed neural activity (Golowasch, Abbott, & Marder, 1999), and respond by inducing restorative changes in ion channel expression (Temporal et al., 2014; Thoby-Brisson & Simmers, 2002) and localization (Mizrahi et al., 2001; Ransdell et al., 2012; Schulz & Lane, 2017). The overall effect, then, is to tune both membrane excitability and synaptic connectivity to moderate neural activity.

Crucially, it is not just neural activity *levels* that are reliably restored after perturbation of the pyloric circuit. Neurons are restored to bursting in their functionally appropriate order—LP, PY, PD. The pyloric CPG, unlike *in vitro* or cortical preparations, provides evidence that ADHP can reliably maintain higher order properties through local activity feedback control. But how exactly is this possible? According to current hypotheses, ADHP mechanisms have no means of detecting burst sequence or phase, only individual neural activity levels. Whether pyloric neurons burst in the correct order or with a significant phase-shift, individual neural activity levels would be the same. So why doesn’t ADHP ever mistakenly restore the circuit to an activity-equivalent, but mis-ordered configuration? How can circuit-level properties like burst order be reliably maintained by regulatory mechanisms operating at the single-neuron level?

At the heart of this conundrum is the fact that circuits with different sets of properties can generate similar network activity, a phenomenon known as parameter degeneracy (Beer et al., 1999; Prinz, 2017) which has theoretical and practical implications. Not only are there many ways to configure a functional pyloric circuit in theory, but experimentally observed conductance densities in identified pyloric neurons are actually highly variable across individuals (Goaillard & Marder, 2021; Goaillard et al., 2009; Schulz et al., 2006, 2007). Recovered circuits, moreover, often exhibit different sets of conductances than they had before perturbation (Alonso et al., 2023; He et al., 2020; Nahar et al., 2012; Rue et al., 2022). ADHP does not simply enforce prespecified values for neural parameters, but acts as a feedback control mechanism allowing parameters to self-organize (Marder & Goaillard, 2006; O’Leary & Wyllie, 2011). The question is, how reliable can we expect this self-organization to be if detected neural activity itself is an unreliable indicator of crucial circuit properties?

Computational models have taken steps toward addressing this question. In conductance-based neuron models, simple regulatory mechanisms have been demonstrated to maintain and restore properties such as firing frequency (Abbott & LeMasson, 1993) and bursting (LeMasson et al., 1993), or even more abstract features like spike adaptation and response to an inhibitory pulse (O’Leary et al., 2014). In general, these model ADHP mechanisms operate by comparing the intracellular calcium concentration, an indicator of average membrane voltage, to a prescribed target value and regulating depolarizing and hyperpolarizing conductances to approach the target. Some such mechanisms have even been able to restore pyloric burst ordering in a model of the pyloric circuit (Golowasch, Casey, et al., 1999).

However, regulatory success in these models is limited to specific hand-picked circuit configurations and perturbations. As described above, this is because averaged neural activity levels are imperfect indicators for such properties of interest. Even when ADHP produces convergence of conductance parameters, homeostatic steady states only sometimes correspond to the correct neural dynamics. To improve reliability in the restoration of bursting dynamics, (Liu et al., 1998) proposed a more complex model calcium sensor made up of three components, each sensitive to calcium fluctuations on a different timescale. They found this sensor improved regulatory performance because its components could more reliably distinguish bursting, which requires dynamics on multiple timescales, from other profiles like tonic firing (Liu et al., 1998). But even this approach failed to generate perfect regulatory performance across all possible initial conditions (Alonso et al., 2023). Analyzing the operation of this compound sensor on a database of model pyloric networks, (Günay and Prinz, 2010) found that functionally ordered networks were likewise not completely distinguishable from dysfunctional ones, no matter how precisely sensor settings were optimized (Günay & Prinz, 2010).

Faced with the prospect of devising and optimizing ever more complicated activity sensors to detect and maintain all the higher-order properties a neural circuit requires, we might instead ask the question: how good is “good enough” for biology? Real organisms, after all, are not perfectly resilient to all possible challenges. Many diseases of the nervous system have been linked to failures of homeostasis (McGregor et al., 2024; O’Leary, 2018; O’Leary et al., 2014; Vislay et al., 2013; Yang & Prescott, 2023). Evolution, it seems, has produced homeostatic plasticity mechanisms that regulate neural properties well enough, given the perturbations that an organism is likely to experience in its lifetime. Therefore, we can learn a lot by taking a perspective that explains the tradeoffs and limits of homeostasis, as well as its remarkable abilities.

This perspective requires consideration of the relationship between parameter degeneracy, ADHP mechanisms, and the perturbations they remedy. In this study, we optimize and analyze an ensemble of models where a simple ADHP mechanism regulates the higher-order property of pyloricness in a neural circuit. We consider the interaction between features of ADHP, local parameter perturbations, and global parameter degeneracy to provide a principled answer to the question: when can activity-dependent homeostatic plasticity maintain higher-order dynamic properties using local activity information? We extract from our model a general criterion for homeostatic success which clarifies the factors that contribute to circuit robustness, and simultaneously identifies its limits.

## 2 Methods

The base of our model is a three-neuron continuous-time recurrent neural network (CTRNN) capable of fixed and oscillatory dynamics, depending on the choice of network parameters (Beer, 1995, 2006). We adapt experimentally-derived criteria previously used to adjudicate the pyloric character of a spiking neural network (Prinz et al., 2004) to this continuously-valued network model. Based on these criteria, we use an evolution-inspired optimization algorithm (Harvey, 2011) to produce a population of unique networks with pyloric-like dynamics. A subset of network parameters are then perturbed and placed under homeostatic control. That is, neural activity can be used to tune these parameters according to a class of ADHP rules previously laid out for this type of network (Williams, 2005). Crucially, we allow the specifics of these homeostatic tuning rules to vary, testing a variety of ADHP mechanisms within this broad class for their ability to maintain the pyloric character of model circuits.

### 2.1 Defining pyloric rhythms in a CTRNN

We aimed to parameterize model circuits whose neural activations autonomously oscillate in an ordered pattern similar to the pyloric CPG. As such, we label each of the three neurons in our circuit after a constituent neuron type of the canonical pyloric rhythm: the pyloric dilator (PD), lateral pyloric (LP) and pyloric (PY) neurons. Note that the synchronized AB/PD pacemaker kernel will be holistically represented by the PD neuron, and the population of PY neurons is represented as a unified element. The evolution of the network’s state can be fully described by three differential equations:

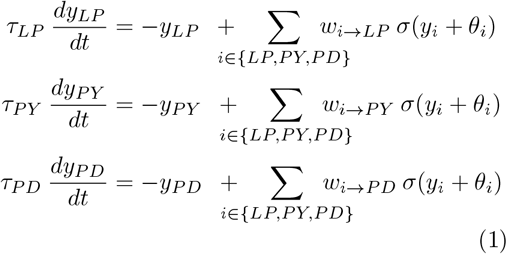

where *y*_*i*_ represents the state of neuron *i* ∈ {*LP, PY, PD*}, *τ*_*i*_ is its time constant, *θ*_*i*_ is the bias of its output function, and *w*_*i*→*j*_ is the synaptic weight connecting unit *i* to unit *j*. The output of neuron *i* is given by *σ*(*y*_*i*_ + *θ*_*i*_), where *σ* is the sigmoid activation function *σ*(*x*) = 1*/*(1 + *e*^−*x*^) (see Figure 1B). These outputs, which are bounded between 0 and 1, may be taken to represent instantaneous neuronal firing rates or proportion of maximum firing rate. Further, we may choose a burst threshold, where time points with an output above this threshold comprise a burst, and those below comprise inter-burst intervals. For the purposes of this study, we set the burst threshold to 0.5, where the activation curve has the steepest slope. Given this threshold, we can parse a time-series of CTRNN node outputs which fluctuate around 0.5 into burst epochs with precise start and end times and inter-burst periods. Neurons whose outputs do not cross 0.5 are effectively considered either silent (below threshold) or tonically firing (above threshold).

**Fig. 1.**
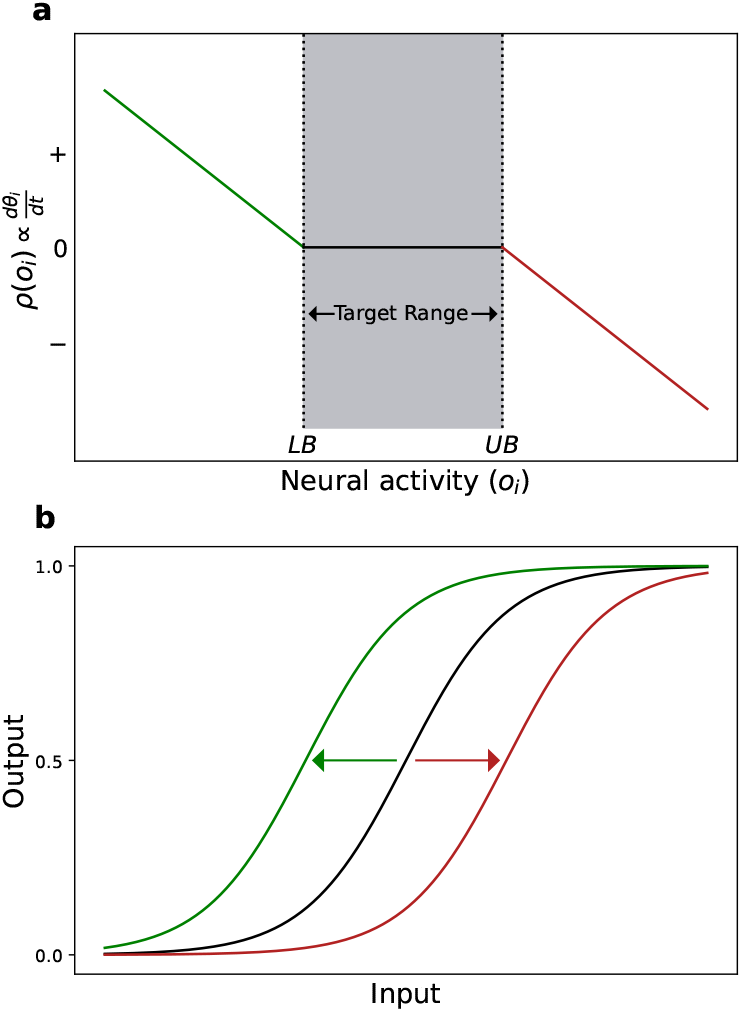
**a**) Plot of generalized homeostatic activation function *ρ* for a given neuron *i* with bias *θ*_*i*_, as a function of neural activity, *o*_*i*_. The rate of change of *θ*_*i*_ is proportional to *ρ. LB* and *UB* denote the lower and upper bounds of the target activity range, within which the rate of change of *θ*_*i*_ is zero. When activity falls below the target range, *ρ >* 0 (**a**, green branch), and *θ*_*i*_ is upregulated, shifting neuron *i*’s activation function, *σ*(*y*_*i*_ + *θ*_*i*_), to the left (**b**, green curve). When neuron *i*’s activation rises above the target range, *ρ <* 0 (**a**, red), so *θ*_*i*_ is downregulated, and the activation function is shifted right (**b**, red).

Using this definition of bursting, we can define criteria for similarity to *in vivo* recordings of the pyloric rhythm. The burst order of the rhythm’s constituent neural populations is LP–PY–PD and the end of the LP burst reliably overlaps with PY. These criteria have been used across models (Günay & Prinz, 2010; Prinz et al., 2004) and experiments (Hooper, 1997) to evaluate and characterize the pyloric pattern. They are not, however, sufficiently determined to apply generically to any repeating rhythm, so we must further require that the LP burst begins during a period of silence from the other two neurons. This assumption agrees well with most *in vivo* pyloric recordings (Hartline & Gassie, 1979; Hooper & Marder, 1987; Rezer & Moulins, 1983) and allows the beginning of the LP burst to serve as the rhythm’s anchor—a starting point from which we can also guarantee that each neuron’s burst begins then ends within one cycle of any valid rhythm. In sum, we extract three binary ordering criteria which our model rhythms must simultaneously satisfy to be considered pyloric:

- All neurons silent before *LP*_*start*_
- *PY*_*start*_ *< LP*_*end*_
- *LP*_*end*_ *< PY*_*end*_

If a given rhythm contains exactly one unbroken burst from each neuron per cycle, there are 5! = 120 different ways to order the beginning and end of each burst, resulting in 120 qualitatively different burst order possibilities. Of these, 10 meet our three criteria for pyloricness (see Figure 10 in the Appendix). Some researchers have further restricted this space of possibilities, assuming that the order of events in the rhythm conforms exactly to: *LP*_*start*_, *PY*_*start*_, *LP*_*end*_, *PY*_*end*_, *PD*_*start*_, *PD*_*end*_ (Hooper, 1997). In light of inter-individual and inter-species variability, we elected to begin our investigation with a more permissive set of requirements based on minimum consensus criteria.

### 2.2 ADHP in a CTRNN

Activity-dependent homeostatic plasticity can be readily implemented in this network model. Generically, ADHP transforms network parameters (in this case, neuron biases, *θ*_*i*_ and connection weights, *w*_*ij*_) into state variables whose rate of change is some function of neural activity and/or activity history. This function usually invokes the relationship between current neural activity and a “set point” or “target”, at which the rate of change in parameters would be zero by definition (O’Leary & Wyllie, 2011). Further, it is typically assumed that the timescale of neural activity is significantly faster than that of ADHP, reflecting assumptions about its molecular mechanisms and allowing a partition of these models into slow and fast subsystems. Experimenters may then justifiably describe neural dynamics as if homeostatically controlled parameters were constant over short intervals and describe parameter dynamics as if neural dynamics were instantaneously equilibrated over long intervals (Rinzel, 1987). This convenient approximation may require greater timescale separation than is often employed or assumed, (Stolting et al., 2023) but can be quite fruitful for analysis. Finally, though there are many documented ways that neurons in a network can persistently influence each other’s electrical properties (i.e. glial signaling, secreted factors, retrograde regulation; Maffei & Fontanini, 2009; Turrigiano, 2012), it is typically assumed that a neuron’s own activity is the only control signal used by ADHP to tune its excitability and synaptic connections. This makes sense from a control-theory perspective, as a neuron’s own activity is the most direct product of these properties. For our ADHP model, we adopt these conventions.

The specific form ADHP takes in our model is based on that devised for CTRNNs by Williams (2004, 2005). As described above, placing a parameter under homeostatic control turns it into a variable whose rate of change is a function of the activity of the neuron to which it is associated. According to the locality assumption, a neuron’s own activity is used to regulate its bias and incoming synaptic weights. For the remainder of this study, however, we will restrict homeostatic regulation to biases, noting that modulating a neuron’s bias is mathematically equivalent to coordinately modulating all its incoming weights. Several experiments have also demonstrated that the intrinsic properties of pyloric neurons have at least as much influence on their relative phase as synaptic properties (Rabbah & Nadim, 2005) and that their regulation is more fundamental to homeostatic rhythm recovery (Thoby-Brisson & Simmers, 2002). Furthermore, to serve intuition-building visualizations of our model, we chose to initially restrict homeostatic regulation to two of the three neuronal biases. We arbitrarily select *θ*_*LP*_ and *θ*_*PD*_. ADHP is therefore described by the following differential equations:

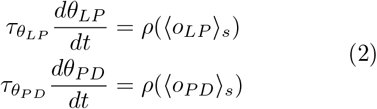

where 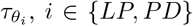 is the time constant of *θ*_*i*_’s change and *o*_*i*_ is the relevant neuron’s time varying output. The value ⟨*o*_*i*_⟩ _*s*_ is the simple moving average, or sliding window average, of neuron *i*’s output, with window length *s* seconds, such that 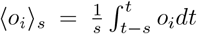 at time *t*. Finally, *ρ*(*x*) is the piecewise continuous homeostatic activation function,

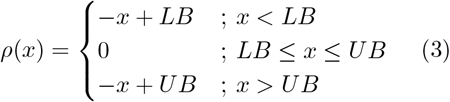

where *LB* and *UB* are the lower and upper bounds of the target activity range (Fig 1a).

All in all, ADHP in our model is a relatively simple controller. When the detected activity of a neuron falls below its target range, its bias is increased, shifting its activation function to the left and increasing the output for each value of its internal state. Conversely, when activity goes above the target range, the neural bias is decreased, shifting the activation function right and decreasing output (Figure 1B). The speed of these processes relative to the rate of change of neural activity fluctuations is slow, enforced by the requirement that 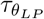 and 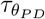 are at least fifty times larger than all neural time constants. Additionally, to reflect the fact that membrane conductances are not infinitely scalable and have physical limitations, we enforce boundaries beyond which neural biases cannot grow or shrink. For the following simulations, biases are restricted to the range [−20, 20].

It has been shown that application of this mechanism with central target ranges (*LB* = .25, *UB* = .75) increases the propensity for oscillations in CTRNNs (Stolting et al., 2023; Williams, 2005). This is because the targets are best satisfied when each neuron’s activation function *σ*(*θ*_*i*_ + *y*_*i*_) is centered over the range of inputs it receives. Circuit configurations where this applies to all neurons, known as center-crossing configurations, are disproportionately more likely to exhibit limit cycles than non-center-crossing networks because neurons have maximal input sensitivity and more readily participate in oscillatory dynamics (Mathayomchan & Beer, 2002). Thus, in pushing circuits towards center crossing configurations, ADHP is systematically encourages oscillation. However, there is no reason to suspect that these oscillations should systematically favor any specific order (i.e. pyloric ordering), much less the finer details of relative phase and timing which may be important for circuit function. The question remains, then, under what conditions can this activity-dependent homeostatic plasticity mechanism successfully regulate the circuit-level property of “pyloricness” using only local activity information.

## 3 Results

### 3.1 Optimization of pyloric networks

Our first step toward enumerating regulatory outcomes in this model was to generate a population of differently parameterized CTRNNs which meet circuit-level criteria for pyloricness. Genetic algorithms are particularly well-suited to this type of ensemble modeling optimization. Their stochastic nature ensures a differently parameterized solution every time the algorithm is run, exploring the space of possibilities with minimal prior assumptions (Beer, 1997; Beer & Gallagher, 1992). Similar techniques have even been used to parameterize conductance-based models of pyloric cells and networks (Alonso & Marder, 2019, 2020; Smolinski & Prinz, 2009).

For our case, the non-homeostatic three-neuron network described in section 2.1 has 15 free parameters: three neural time constants, three neural biases, and nine connection weights. Thus, a vector of values for these 15 parameters completely specifies a CTRNN, which is numerically simulated and evaluated for its pyloricness according to a defined fitness function. Fitness, more specifically, is awarded to a circuit in all-or-nothing steps for each pyloric requirement it satisfies. First, for each of the three neurons whose output crosses above and below the burst threshold, we add 0.05 points. Then, if all three neurons have defined bursts, the circuit is evaluated on the three independent ordering criteria listed in section 2.1; 0.05 points are added if the LP neuron begins its burst during silence from PY and PD, an additional 0.05 if the PY neuron begins its burst before LP’s burst ends, and a final 0.05 if the LP neuron ends its burst before PY does. Any circuit that meets all six of these criteria, accruing a base fitness of 0.3, is considered pyloric.

From a randomly parameterized initial population of 100 circuits, individuals with the highest fitness are copied and randomly mutated, while those with the lowest fitness are removed from the population (Harvey, 2011). After 100 iterations of this process, the best circuit is likely to produce a pyloric oscillation. If it does, its parameter set is collected in a database. We aimed to produce two databases of 100 pyloric circuits each: one where correct burst order was the only requirement for inclusion and one where realistic phase and timing of bursts was also required. To fill the first database, we simply repeated the evolution procedure until we had evolved 100 circuits that met all pyloric burst order criteria. For the second database, we prescribed a way for pyloric circuits to earn additional fitness if the relative phasing of their bursts approximated experimentally-observed timing characteristics from a set of lobster (*H. americanus*) preparations (Prinz et al., 2004). Specifically, the z-scores (calculated with experimentally reported averages and standard deviations) for each neuron’s duty cycle (proportion of the cycle spent bursting), as well as for the phase delay from the start of the PD burst to the start of the LP and PY bursts, were averaged, inverted, and added to fitness. To be entered in this more restrictive database, circuits had to attain a minimum fitness of 2, with 0.3 awarded for meeting pyloric ordering criteria and at least 1.7 awarded for timing similarity.

While they all meet criteria for pyloricness, rhythms produced by circuits in the first database had some qualitative differences from each other. Of the ten distinct rhythm classes which meet our criteria for pyloricness (see Figure 10), eight are represented by circuits in our first database. In the second database, where a timing similarity requirement was imposed, all circuits were of the same type— [*LP*_*start*_, *PY*_*start*_, *LP*_*end*_, *PY*_*end*_, *PD*_*start*_, *PD*_*end*_]. This is the ordering which would necessarily result from perfect conformity to experimental means on all five relative timing metrics, and is commonly observed in STG preparations (Hooper, 1997). Later (section 3.4), we will utilize the first dataset to examine the consequences of increased heterogeneity for homeostatic generalizability. For now, however, we will focus on this second, more homogeneous database of pyloric circuits.

Despite their dynamic similarity, circuits in this database exhibited significant parameter variability. Seven of the 15 network parameters spanned *>*94% of their allowed range. Unfortunately, our sample is too small to support meaningful analysis of the structure of this variability (i.e. linear and nonlinear relationships between parameters of evolved circuits, the geometric organization of pyloric CTRNN solution space). We leave this for future work and refer the reader to other ensemble modeling contexts where such analyses which paint vivid pictures of the parametric landscapes on which ADHP must operate (Ball et al., 2010; Beer, 2006; Doloc-Mihu & Calabrese, 2011, 2014; Foster et al., 1993; Goaillard & Marder, 2021; Hudson & Prinz, 2010; Lamb & Calabrese, 2013; Mittal & Narayanan, 2018; Sutulovic et al., 2025; Taylor et al., 2006, 2009). However, it suffices for the current study that the parametric variability present among our evolved circuits is consistent with variable conductance densities in real CPGs (Calabrese et al., 2011; Marder et al., 2007; Schulz et al., 2006) and that it allows us to assess the varied conditions under which ADHP can or cannot maintain pyloricness against circuit perturbations.

Importantly, by virtue of their differing parameter sets, the circuits in our dataset respond differently to equivalent parametric perturbations. Perturbing the parameters of evolved circuits can produce a range of results, from deforming the shape and timing of neural bursts without violating pyloric criteria, to changing the relative position of rhythm components such that pyloric criteria are violated, to disrupting the bursting of one or more neurons such that pyloric criteria are undefined. In subsequent analyses, we will focus on the effects of perturbing the two parameters which will eventually be placed under homeostatic control: *θ*_*LP*_ and *θ*_*PD*_. The response of three example circuits to a range of perturbations of these bias values are plotted in Figure 2.

**Fig. 2.**
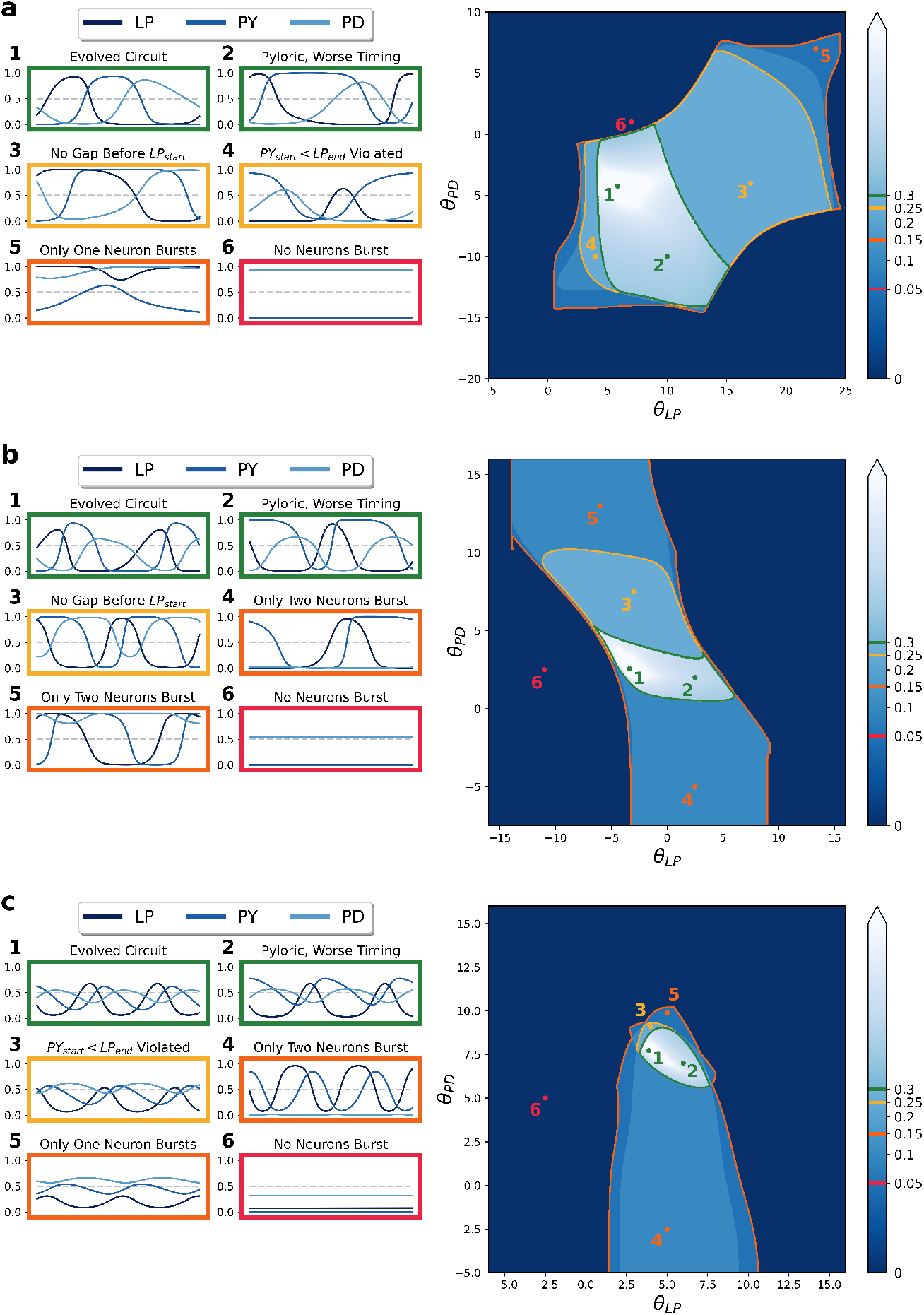
Slices of CTRNN parameter space which cut through three unique evolved pyloric circuits. Each large plot demonstrates the effect of varying *θ*_*LP*_ and *θ*_*PD*_ while keeping other parameters constant at evolved values, with our pyloricness metric plotted in shades of blue. Colored lines demarcate regions of qualitatively different steady-state behavior, evaluated after a transient of 50 seconds (green-pyloric ordering criteria met, yellow-subset of ordering criteria violated, orange-one or more neurons not bursting, outside-no neurons bursting), with example neural output time series from representative circuits in each region (marked by numbered points).

Each subfigure depicts a two-dimensional slice through CTRNN parameter space, over which *θ*_*LP*_ and *θ*_*PD*_ vary while all other parameters are held constant at the values of an evolved circuit. We outline several contiguous regions where equilibrated neural activity is similar with respect to our pyloric criteria, and plot neural output time-series for representative circuits in each region. Boundaries between regions mark sudden changes in pyloricness, such as where limit cycles no longer satisfy one or more ordering criteria or certain neurons no longer cross the bursting threshold. The outermost boundary, where all neurons stop bursting, often roughly corresponds with a limit-cycle-terminating bifurcation. The correspondence is only approximate because usually, as in the case of a supercritical Hopf bifurcation, the limit cycle smoothly decreases in size prior to disappearance, and this shrinkage prevents neural states from crossing the burst threshold even while they still oscillate (see for example Fig 2a, timeseries 5). Finally, while we did not denote it in these plots, a small subset of perturbed circuits are multistable, where more than one stable equilibrium co-exists in neural state space. Thus, a single parameterized circuit may have different pyloric status depending on the steady state to which it equilibrates. Multistability, however, was rare in the relevant slices through parameter space, so each parameter set can generally be assumed to correspond to one stable limit set with definitive pyloric or non-pyloric status.

Thus, we have produced a database of 100 parametrically distinct CTRNNs which generate similar pyloric patterns. Perturbing the parameters of these networks, in imitation of a bodily or environmental perturbation to the STG, produces different results depending on the perturbation and the circuit’s unperturbed parameters. The perturbations we will consider are translations within the two-dimensional slices of parameter space centered on evolved pyloric solutions and extended across ADHP-controlled parameters, *θ*_*LP*_ and *θ*_*PD*_. A useful ADHP mechanism, then, should act upon these same parameters to restore pyloricness after a variety of perturbations.

### 3.2 Optimization of homeostatic mechanisms

We next asked whether activity-dependent homeostatic plasticity acting concurrently upon the biases of LP and PD could restore pyloricness following perturbations to those parameters. To optimize ADHP mechanisms for this task, we used another genetic algorithm with the same overall structure as in the previous section. A single ADHP mechanism is described by four “meta-parameters” per bias parameter, *θ*_*i*_ that it regulates: the lower bound of its target range (*LB*_*i*_), target range width (*UB*_*i*_ − *LB*_*i*_ = Δ_*i*_), sliding window size (*s*_*i*_), and time constant of regulation 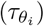. In our case where ADHP regulates *θ*_*LP*_ and *θ*_*PD*_, this makes eight ADHP meta-parameters. During evolution, *LB*_*i*_ was allowed to assume values anywhere in the possible output range of a neuron, from 0 to 1. The width of the target range could also be any value between 0 and 1, with the upper bound *UB*_*i*_ clipped to 1 if necessary. The sliding window duration (in seconds) was restricted to the interval [0,10], with a choice of 0 corresponding to instantaneous activity detection. The time constants of regulation were restricted to the interval [100, 200]. This ensured that regulation was slow compared to the dynamics of neural states, whose time constants were restricted to the range [0.1, 2], resulting in a minimum timescale ratio of 50:1. This is a much stricter timescale separation than in previous implementations of this ADHP model, where the dynamics of the slow subsystem frequently generate oscillations on behavioral timescales (Stolting et al., 2023; Williams, 2004).

To evaluate a single ADHP mechanism for regulation of a given pyloric network, we first perturb the network’s LP and PD biases to a value on an evenly-spaced 3-by-3 grid of points ({−9, 0, 9}^2^), allow the network to equilibrate for 250 seconds, and then activate ADHP and allow it to regulate parameters for 2500 seconds before evaluating the network’s pyloric fitness. To discourage overreliance on the exact duration of regulation, pyloric fitness is re-evaluated after another 2500 seconds, and the two fitness values are averaged together. This result is averaged across trials starting from all nine initial conditions. Finally, to prioritize ADHP mechanisms that recover pyloricness from more initial points over those that recover it with better timing from fewer initial points, we award a flat fitness bonus for each run where the final circuit meets pyloric criteria.

For each of the 100 pyloric circuits in our database of interest, we evolved five ADHP mechanisms, starting each evolutionary trial with a randomized population of 50 regulators which evolved for 100 generations. For 71 of those circuits, at least one ADHP mechanism evolved to maintain pyloricness against all *θ*_*LP*_ and *θ*_*PD*_ perturbations, even on a more extensive 6-by-6 grid ({−15, −9, −3, 3, 9, 15}^2^). Of the remaining 29 circuits, 26 had at least one ADHP mechanism that restored pyloricness after some subset of perturbations, though not all of them. That leaves 3 circuits for which we were unable to evolve any ADHP mechanism to maintain pyloricness at all.

Noting the differential performance among these 500 evolved ADHP mechanisms, we first asked whether the distribution of ADHP metaparameters exhibited any trends that correlated with success or clear evolutionary pressure (Figure 3). The lower bounds of the target ranges, *LB*_*LP*_ and *LB*_*PD*_, were unimodally distributed with peaks at intermediate activity values (**a**,**e**). While there were no robust differences in distribution between successful, mixed, and unsuccessful ADHP mechanisms, this reflects meaningful regulation requirements that we will be able to clarify later on. Next, the width of the target activity ranges (Δ_*i*_) was low in almost every evolved mechanism (**b**,**f**), and this trend was even more striking for successful regulators. In the case of Δ_*PD*_, no successful regulators had target ranges wider than 0.373 (**f**). We note this trend for further analysis.

**Fig. 3.**
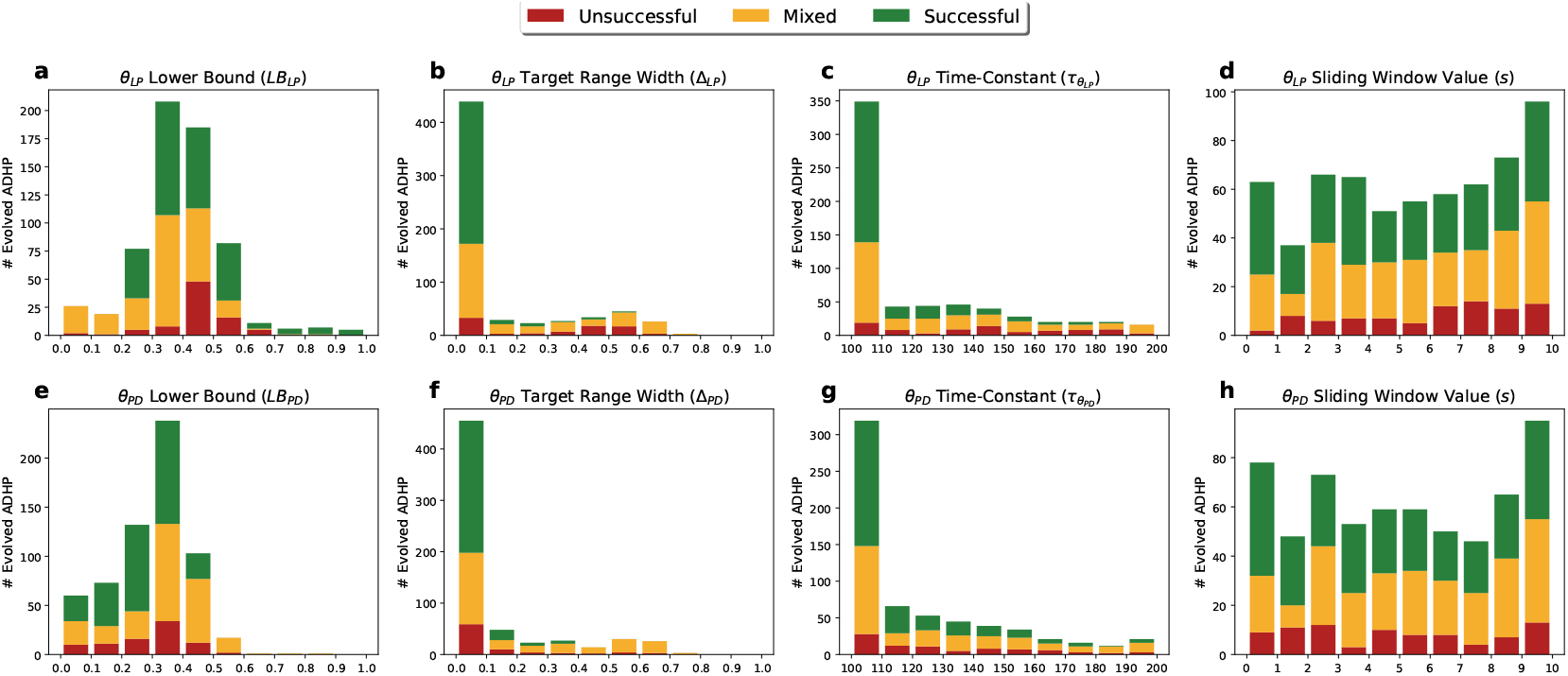
Distribution of evolved ADHP meta-parameters across allowed ranges (*x*-axes) for mechanisms with no success (red), mixed success (yellow), or full success (green) in restoring pyloricness after multiple perturbations. Five ADHP mechanisms were optimized for performance on each of 100 pyloric circuits. Distributions for the lower bounds of the target ranges (*LB*_*i*_) appear unimodal with peaks below 0.5 (**a**,**e**). The widths of evolved target ranges (Δ_*i*_) were strongly selected to be small, and in the case of *θ*_*PD*_, no successful regulators had range widths above 0.373 (**b**,**f**). Time constants of regulation 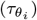 were also selected to be small, but successful regulators are found throughout the allowed range (**c**,**g**). The length of sliding average windows (s) were uniformly distributed with no obvious trends (**d**,**h**).

Evolved time constants of regulation 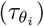 were also concentrated at the lower end of their range (**c**,**g**). However, we hypothesized that this trend was a result of artificial pressure imposed by our choice of simulation time. To test this hypothesis, we reevaluated a randomly selected subset of evolved ADHP mechanisms, but doubled their 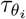 values and doubled the allowed simulation time. Each mechanism fell into the same performance category as before. The most likely explanation for this trend, then, is that the shorter simulation time was inadequate for slower ADHP mechanisms to reach their targets. Later, we will mathematically demonstrate why the time constant of regulation does not affect the outcome of ADHP, only how quickly it is reached.

Lastly, there were no obvious trends in sliding window durations, which were relatively uniformly distributed throughout the allowed range across performance levels (**d**,**h**). We therefore highlight only the most robust correlates of regulatory success: low-to-intermediate activity targets and narrow target ranges. Through subsequent analyses, we will be able to explain why these features are beneficial. Next, we took a closer look at the parameter trajectories induced by evolved ADHP mechanisms. Representative examples in Figure 4 illustrate that regulated parameters *θ*_*LP*_ and *θ*_*PD*_ tend to converge on one or multiple endpoints in parameter space. When trajectories terminate in the pyloric region of the *θ*_*LP*_ *θ*_*PD*_-plane, this means that pyloricness has been restored. As noted above, many evolved circuit-ADHP pairs had this as their only outcome, implying a single pyloric attractor to which all perturbed circuits recover (e.g. Fig 4, left column).

**Fig. 4.**
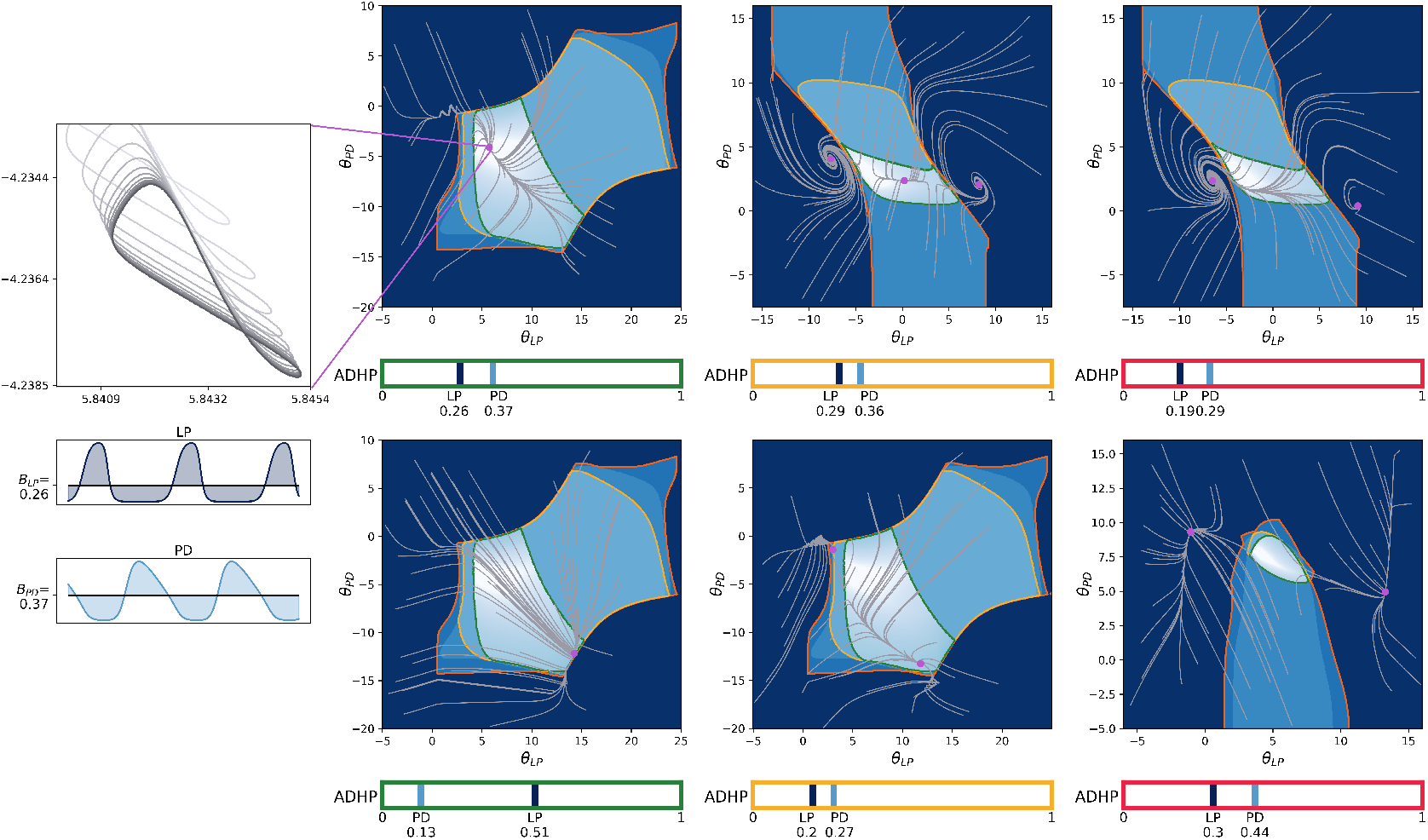
Evolved regulatory mechanisms attract parameter space trajectories to one or several endpoints for reliable, mixed, or unsuccessful recovery outcomes. Each plot considers an evolved ADHP mechanism with different neural activity targets, all of which have ranges with near-zero width (targets at light and dark blue bars), acting upon example circuits whose *θ*_*LP*_ and *θ*_*PD*_ were randomly perturbed from their evolved values. Light gray trajectories track parameter changes over a period of 50,000 seconds (≈13.9 hrs) and purple points mark their values at the end of the simulation. **Left column:** Two reliable ADHP mechanisms responding to perturbations of the first example circuit (see Fig 2a) can recover pyloricness from all perturbations. **Middle:** Mechanisms with mixed success attract some trajectories to the pyloric region and others outside it in the first and second example circuits (see Fig 2 a&b). **Right:** Unsuccessful mechanisms attract all trajectories outside the pyloric region in the second and third example circuits (Fig2 b&c). **Inset:** Zoom in on the approach to a homeostatic “endpoint”, which is actually a small-amplitude limit cycle projected on the *θ*_*LP*_ *θ*_*PD*_-plane. Color saturation increases as the simulation continues. Timeseries below show the outputs of LP and PD neurons in the last 15 seconds of simulation time, as they oscillate around their activity targets (shaded area).

Other ADHP mechanisms did not have such reliably successful outcomes. At the other extreme, if attractors exist only outside the pyloric region, then ADHP cannot maintain pyloricness and in fact actively disrupts it (Figure 4 right column). Even if trajectories happen to pass through the pyloric region, with neurons temporarily exhibiting the correct oscillatory pattern, they continue onward to a dysfunctional regime. This mirrors findings from other models where even logically-structured homeostatic mechanisms can disrupt circuit function (O’Leary, 2018; O’Leary et al., 2014). Finally, in cases where there are multiple regulatory endpoints, some subset of them may be within the pyloric region and others outside it. This makes outcomes perturbation-dependent. Some perturbations leave circuits in the basin of attraction of the pyloric endpoint, and are thus recoverable. Other perturbations place circuits in the basin of attraction of a nonpyloric endpoint, and are thus unrecoverable. Interestingly, this ill-fated basin may include sections of the pyloric region itself.

Though approaching an intermediate endpoint was by far the most common fate for regulated circuits in our dataset, there were occasionally other outcomes. First, ADHP did sometimes press parameters against the imposed boundary conditions. In these cases, trajectories do still converge to a point along the boundary, which may or may not lie in the pyloric region, but their convergence is partially dependent on imposed restrictions. Second, circuits may traverse limit cycles through parameter space. These orbits can cross through different regions of the plane, leading the circuit to cycle through multiple pyloric statuses. We will return to these two alternative outcomes at the end of the next section.

Largely, though, homeostatic recovery trajectories tend to approach a set of endpoints, of which all, some, or none lie in the pyloric region. This results in regulation that succeeds for all, some, or no initial conditions. Our model can thus represent such varied phenomena as functional recovery, perturbation dependence, and pathological compensation. Our next objective is to derive heuristics to explain and predict these regulatory outcomes in our model.

To this end, we closely examined the system’s behavior in the vicinity of these ubiquitous attractive points. As is the case in several other models of neural ADHP (Abbott & LeMasson, 1993; Olypher & Prinz, 2010), we found that the parameter-space equilibrium points of successful ADHP mechanisms are not actually *points* at all, but small-amplitude limit cycles. These limit cycles appear static at reasonable plotting scales (Fig 4 inset), yet evade simple analytical identification. In hindsight, this parameter oscillation seems inevitable, as the only way that neural biases could remain static (*dθ*_*i*_*/dt* = 0) while neural outputs oscillate in a pyloric pattern is if those oscillations were entirely contained in their respective target activity ranges. Given the narrow ranges of our evolved ADHP mechanisms, this is unlikely. The small amplitude of the parametric oscillation results from the timescale separation between the rates of change in neural states and biases, which ensures that brief excursions from the target activity range do not drastically affect bias values, except when changes aggregate over successive cycles. Homeostatic attractors, then, result when excursions above and below the target range cancel out over the course of a limit cycle. Additionally, timescale separation ensures that, for circuits near the homeostatic attractor, neural dynamics look quite similar whether parameters are frozen or continually regulated. Therefore, information gathered about neural behavior occurring with fixed parameters (i.e. parameter space regions demarcated in Figure 2) is reasonably informative about neural behavior when a homeostatic system is passing through that point in parameter space. The fact remains that ADHP attractors cannot just be identified analytically by setting the rates of change equal to zero and solving for system states, but armed with these two observations (balance over the target activity range and timescale separation), we can approximate ADHP-induced flow over parameter space and predict the location of attractors.

### 3.3 Extracting alignment criteria that predict performance

In order to explain the differential success of regulators in our dataset, we must describe the relationship between an ADHP mechanism’s features and its quasi-steady state parameter values. To this end, we will characterize a flow field over the *θ*_*LP*_ − *θ*_*PD*_ plane and use its nullclines to predict homeostatic endpoints and their basins of attraction. Based on the pyloric status of these endpoints, we can predict the ADHP mechanism’s success in restoring pyloricness after a range of parametric perturbations. Then, considering the set of all possible ADHP mechanisms that could regulate a given circuit, we can extract necessary and sufficient conditions for successful regulation based only on the relationship between neural activity and pyloricness in the circuit’s vicinity.

Our characterization of the flow field over parameter space begins with the homeostatic equations which define the rate of change in network parameters (Equation 2). From there, we invoke the assumption that the slow timescale of ADHP is well separated from that of the neural states (with time constants 100-500 times as large). As a result, we assume that as *θ*_*LP*_ and *θ*_*PD*_ change, neural states are instantly equilibrated to some member of the limit set that exists for the non-homeostatic CTRNN with those parameters. Additionally, as we observed in the previous section, the homeostatically-induced rate of change in parameters is slow enough relative to neural states that it only produces a significant effect in the aggregate over time. In the case of oscillatory neural activity, we will be interested in the average homeostatic rate of change over one cycle of the neural states. This strategy of approximating the rate of change of the slow subsystem by averaging its value over one cycle of the fast subsystem has been fruitfully applied in other neural models, as well (Abbott & LeMasson, 1993; Lemaire et al., 2025; Olypher & Prinz, 2010; Rinzel, 1987). Finally, for any circuit with *θ*_*LP*_ and *θ*_*PD*_ set to specific values 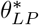 and 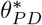, where neural states have reached the *n*th member of the non-homeostatic circuit’s set of limit sets, let 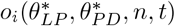 be the function that describes the time-varying output of neuron *i* ∈ {*LP, PD*}.

Altogether, we have

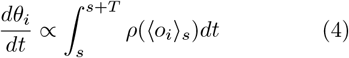

When the asymptotic behavior of the neural states is oscillatory, *T* denotes the period of oscillation. Otherwise, *T* is fixed at an arbitrary value. Note that, if the set of limit sets has only one member at 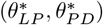, this value is singly defined. Otherwise, if the circuit is multistable, the value depends on which basin of attraction the neural states are in when 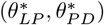 is attained.

Because sliding window averaging did not seem to have much bearing on regulatory success among evolved homeostatic mechanisms, we will simplify our analysis by considering ADHP without it (*s* = 0, ⟨*o*_*i*_⟩ _*s*_ = *o*_*i*_). Further, we will partition the interval [0, *T*] into segments where neural outputs are above, within, or below the target range, [*LB*_*i*_, *UB*_*i*_]. Without changing the value of the integral, we can also reorder these segments such that the interval [0, *T*_1_) captures all values of *o*_*i*_ *< LB*_*i*_, the interval [*T*_1_, *T*_2_) captures values where *LB*_*i*_ *< o*_*i*_ *< UB*_*i*_, and [*T*_2_, *T*] captures values of *o*_*i*_ *> UB*_*i*_. This re-ordered (potentially discontinuous) timeseries of neural outputs will be denoted *ô*_*i*_. Finally, it will be useful for later analysis to express *UB*_*i*_ as the sum of *LB*_*i*_ and the width of the target range, Δ_*i*_. With this, we have

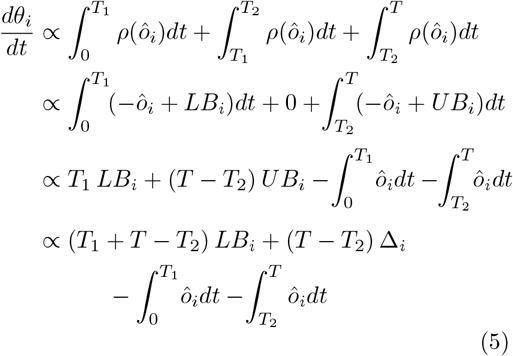

Using this expression, then, we can calculate the sign of the net flow field over parameter space at any point, 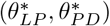. If neural states approach a fixed point at 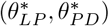, they will remain below, above, or within the target range boundaries (consider *T*_1_ = *T*_2_ = *T, T*_1_ = *T*_2_ = 0, or *T*_1_ = 0∧*T*_2_ = *T*, respectively), so we need only compare the value of the fixed point to the target range. In any case, since we are primarily interested in the homeostatic endpoints where *dθ*_*i*_*/dt* = 0, we can determine from expression 5 whether a point 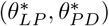 lies on the nullcline (elsewhere called the isoline; Olypher and Prinz, 2010) in the *θ*_*i*_ dimension, for ADHP with a given *LB*_*i*_ and Δ_*i*_. Most points that make up these nullclines are stable and locally attractive, with positive values of *dθ*_*LP*_ */dt* lying to the left of the nullcline and negative values to the right, or in the case of *dθ*_*PD*_*/dt*, positive values below the nullcline and negative values above. Stability occurs more often than instability in this system because increasing the bias value of a CTRNN neuron shifts its activation function to the left, which tends to increase its output (see Figure 1). It is this intuition, after all, that guided the formulation of this ADHP model. We will see, however, that complex network interactions can lead to the opposite arrangement, especially in oscillatory regions of parameter space. In any case, points 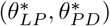 which lie on the stable portion of both nullclines will be quasi-steady states of the homeostatic slow subsystem.

When the width of the target range is zero, as is the case for many of our evolved solutions, expression 5. can be significantly simplified. For this, we define the single target activity value *B*_*i*_ = *LB*_*i*_ = *UB*_*i*_, set Δ_*i*_ = 0, and set *T*_1_ = *T*_2_ to obtain the following expression for the net direction of ADHP regulation with a single target activity value:

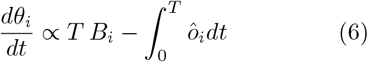

We can further see that, when the width of the target activity range is zero, nullclines for ADHP occur where the following equation is satisfied:

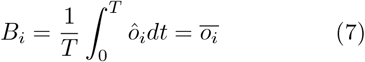

In other words, they occur where the target activity value is equal to the time-averaged value of neural output 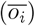. This result is applied in Figure 5 to analyze select circuit/ADHP pairs that we highlighted in Figure 4. For points in the *θ*_*LP*_ *θ*_*PD*_-plane surrounding our three example circuits (see Figures 2 & 4), we calculated the asymptotic average value of LP (Fig 5, top) and PD neurons (Fig 5, bottom), then plotted contour lines where this average is equal to the neuron’s activity target (Fig 5, solid lines) for a specified ADHP mechanism. Solid points where stable portions of these lines cross correspond to homeostatic endpoints from earlier simulations. For some circuit/ADHP pairs, stable crossings exist only within the pyloric region (left column), and this produces reliable regulation across all possible perturbations. For others, they exist both within and outside the pyloric region (center column), producing mixed or perturbation-dependent regulation. And for some, they exist only outside the region (right column) or not at all, producing disruptive regulation. Thus, based only on the asymptotic behavior of fixed, non-homeostatic circuits in the domain of a given ADHP mechanism, we can predict the success of that mechanism in regulating pyloricness.

**Fig. 5.**
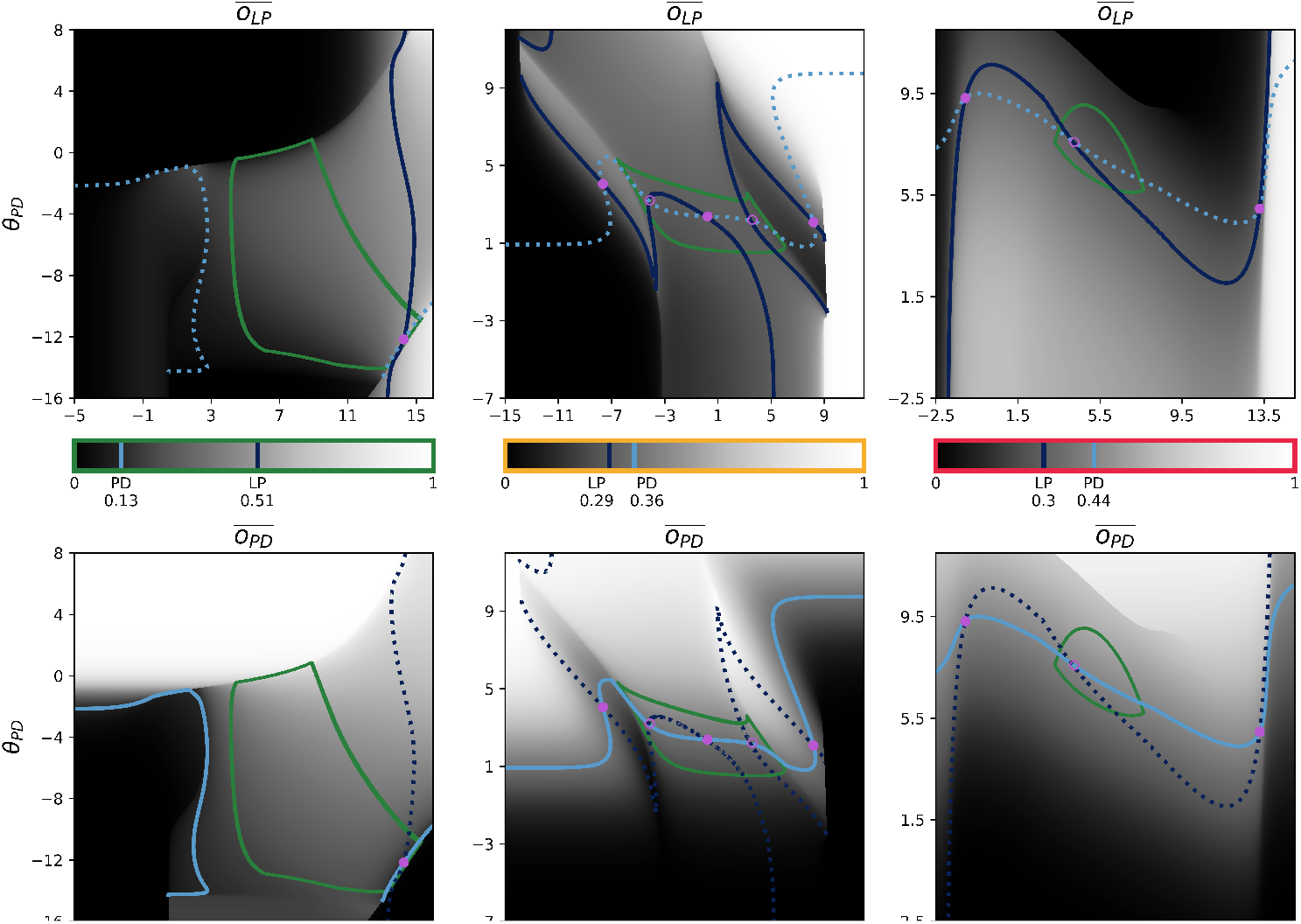
Homeostatic nullclines plotted for select ADHP mechanisms acting nearby example pyloric circuits. The color scale in the center of each column is marked with the activity targets for LP (dark blue) and PD (light blue) and the average value (see expression 7) of the LP neuron (top) and PD neuron (bottom) are plotted in grayscale. Superimposed on these plots are the nullclines where *dθ*_*LP*_ */dt* = 0 (dark blue) and where *dθ*_*PD*_*/dt* = 0 (light blue), where the nullclines aligned with the averages depicted in each plot are solid and the nullcline derived from the other neuron’s average is dashed. Homeostatic endpoints from corresponding simulations (see Fig 4) are marked with solid pink points. Newly identified unstable points are marked with open pink points.

Since any point in the *θ*_*LP*_ *θ*_*PD*_-plane could be made into a homeostatic endpoint by setting the targets of ADHP equal to its neurons’ asymptotic averages, we can analyze the range of regulatory possibilities by examining the distribution of averages and their corresponding pyloric status. More concretely, if we record the asymptotic neural averages 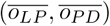 corresponding to each point 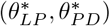 in the pyloric region and the asymptotic neural averages for each point outside the pyloric region, then successful ADHP mechanisms are those that target averages occurring only among pyloric circuits and not elsewhere. Even though individual neural activity does not, in general, specify the circuit-level characteristic of pyloricness, it only matters that, among circuit configurations ADHP can reach, satisfying target activity values happens to imply proper burst order. ADHP mechanisms with mixed success are those which target averages occurring both inside and outside the pyloric region. For these mechanisms, target activity values may sometimes correspond to pyloric burst order and sometimes not; the outcome depends on which basin of attraction the circuit was initially perturbed to. And finally, ADHP that fails to regulate pyloricness may either target values that occur exclusively outside the pyloric region, or it may lack a steady state at all (i.e. the average values ADHP targets do not co-occur in its domain or are unstable).

In Figure 6, we test these intuitions by comparing the value, stability, and pyloricness of co-occurring averages in the vicinity of our three example circuits to the performance of simulated ADHP mechanisms with zero-width target ranges, no sliding window averaging, intermediate time constants 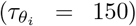, and all possible combinations of targets (*B*_*LP*_, *B*_*PD*_). Up to minor detection errors induced by sampling coarseness and sharp pyloric boundaries, we find that in each of our example circuits, predictions align with simulations.

**Fig. 6.**
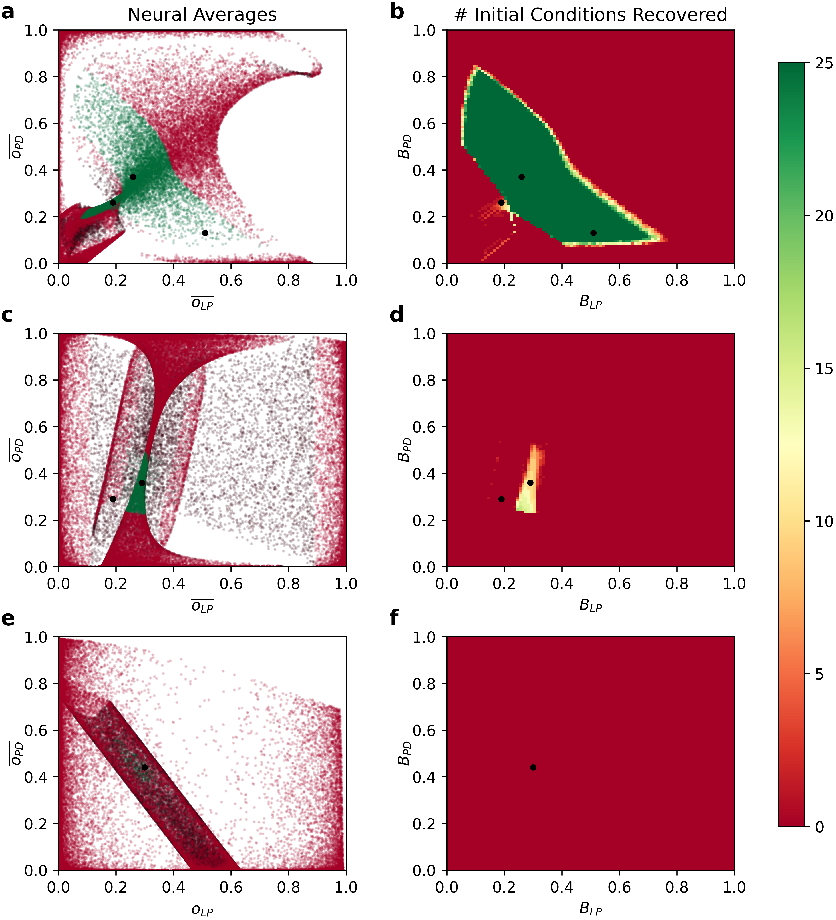
Existence of neural averages among pyloric and non-pyloric circuits predicts performance of ADHP with zero range. In the *θ*_*LP*_ *θ*_*PD*_-plane running through each of three example circuits, we examined 10,240,000 CTRNNs on a grid (*θ*_*LP*_, *θ*_*PD*_ ∈ [−16, 16] *×* [−16, 16]) and recorded the average value of their LP and PD neurons over either one cycle of the rhythm or 50 seconds for non-oscillatory circuits, as well as stability, and pyloric status. Plotted points (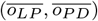) are colored green if pyloric and red if non-pyloric, with brighter versions of these colors indicating stability and dark versions instability. For zero-range ADHP mechanisms with 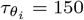 and (*B*_*LP*_, *B*_*PD*_) picked from a grid on the range [0, 1]*×*[0, 1], we simulated recovery of the circuit following 25 perturbations of (*θ*_*LP*_, *θ*_*PD*_) to points on the grid {−10, −5, 0, 5, 10}^2^. Right plots depict the number of times out of 25 that pyloricness was recovered after 50,000 seconds, for each ADHP mechanism. There is strong accordance between left and right plots for all three example circuits, with successful regulation targeting stable average values that occur exclusively in the pyloric region. Successful regulation is possible in our first example circuit (**a**,**b**). Only partial success can be achieved for our second circuit (**c**,**d**), as pyloric averages are inseparable from non-pyloric ones. Regulation is not possible for our third circuit because all pyloric averages are unstable with respect to this regulatory mechanism (**e**,**f**).

More generally, we can explain disparities in regulatability across circuits. Within the set of ADHP-reachable circuit configurations (in this case the *θ*_*LP*_ *θ*_*PD*_-plane which runs through a circuit from our database), there may or may not *exist* a pair of neural averages 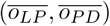 which occurs exclusively in the functional region. Surrounding our second example circuit, for instance, every pair of neural averages that exists in the pyloric region also exists somewhere outside it (Fig 6c). Therefore, the best ADHP can possibly do is to target one of these values and achieve perturbation-dependent success. This is why our optimization algorithm and later systematic search of ADHP meta-parameter space both failed to produce a perfectly reliable regulator. In the case of our third example circuit, none of the neural averages that occur in the pyloric region meet the criteria for regulatory stability. More specifically, 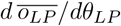 is positive throughout the pyloric region. Increasing the bias of the LP neuron in an attempt to compensate for below-target activity paradoxically lowers average activity levels even more, pushing trajectories out of the pyloric region. As a result, ADHP cannot even partially regulate this circuit (Fig. 6f).

Of the 100 pyloric circuits in our database, 80 can be reliably regulated because there exist pairs of average values which occur only within the pyloric region and nowhere else in the plane of accessible configurations. There were 14 circuits for which perfect regulation was not possible because pyloric averages were inseparable from non-pyloric ones. There was only one circuit (the one we have been examining) for which even mixed regulation was impossible because all pyloric averages were unstable.

Also of note, for each of the 100 pyloric circuits in our data set, the best zero-range ADHP mechanism was always as good as or better than the best evolved ADHP mechanism. This supports our earlier hypothesis that there is nothing to be gained in this particular model from using sliding window averaging, adjusting the time constants of regulation, or widening the activity target range. In fact, we can now explain why broader target ranges only ever hurt performance. Perfect regulation against all perturbations on the *θ*_*LP*_ *θ*_*PD*_-plane can be achieved by placing a single homeostatic endpoint in the pyloric region and excluding all endpoints that would occur outside it. A wider target range cannot produce any more pyloric endpoints than a narrow one. All it can do is capture more static (and therefore non-pyloric) endpoints, if they exist in the plane. Narrow target ranges, in other words, minimize the possibility for ADHP to stabilize “false positives”—circuit configurations that meet activity targets but are not functional. False negatives, on the other hand, where ADHP passes over some functional circuit configurations, are not at all detrimental, but entirely necessary for this kind of regulation. This point has implications for interpreting parameter variability in real neural circuits. Just because multiple circuit configurations may in principle be equally functional, they may not necessarily be observed in a circuit whose homeostatic regulation mechanisms utilize proxies for functionality. Accordingly, any observed inter-individual variability of circuit parameters is a more direct result of equivalence of this proxy than equivalence of function or history.

Finally, our dataset contained several circuits for which simulated ADHP performance deviated from predictions. These cases highlight two important considerations to refine our predictive framework: namely, ADHP’s interaction with bifurcation manifolds and imposed boundary conditions. Regulators of one such anomalous circuit exhibited both phenomena in different regions of its meta-parameter space (Fig 7b). For certain regulators (for example, Fig 7 point 1), ADHP targets co-occur only within the pyloric region of circuit parameter space, which should produce robust regulation. However, a subset of trajectories never make it to the predicted pyloric endpoint, trapped instead in a stable parametric limit cycle. This cycle, with a period of approximately 975 seconds and a path which does not enter the pyloric region, does not produce pyloric neural dynamics. Its location does not coincide with any homeostatic nullclines, being far from areas of parameter space where average neural activity approximates homeostatic targets. Essentially, neural parameters following this cycle flip-flop between the regimes of two fixed points, across a band of bistability in between. When states reach the first fixed point (which at the top-left-most point of the cycle is singly stable), the activity of the LP neuron is below its target, while the activity of the PD neuron is above its target. ADHP pushes circuit parameters down and to the right. This introduces a new stable fixed point, but neural states remain within the basin of attraction of the first, tracking it until it loses stability (lower-right-most point of the cycle). At that point, neural states rush to the remaining stable point, where LP activity is now too high and PD activity is too low. Circuit parameters are pushed up and to the left, neural states tracking this new fixed point across the region of bistability, until only the original fixed point remains and the cycle starts again. Thus, even though target averages are never stably attained at any point during the cycle (and indeed do not coincide with any stable fixed point nearby) it attracts nearby homeostatic trajectories. This phenomenon demonstrates that it is insufficient to examine only the set of averages 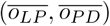 that exist in the plane when predicting regulatory outcomes; we must also interpolate between the values of fixed points which are proximate in parameter space, for their potential to produce anomalous non-pyloric attractors.

**Fig. 7.**
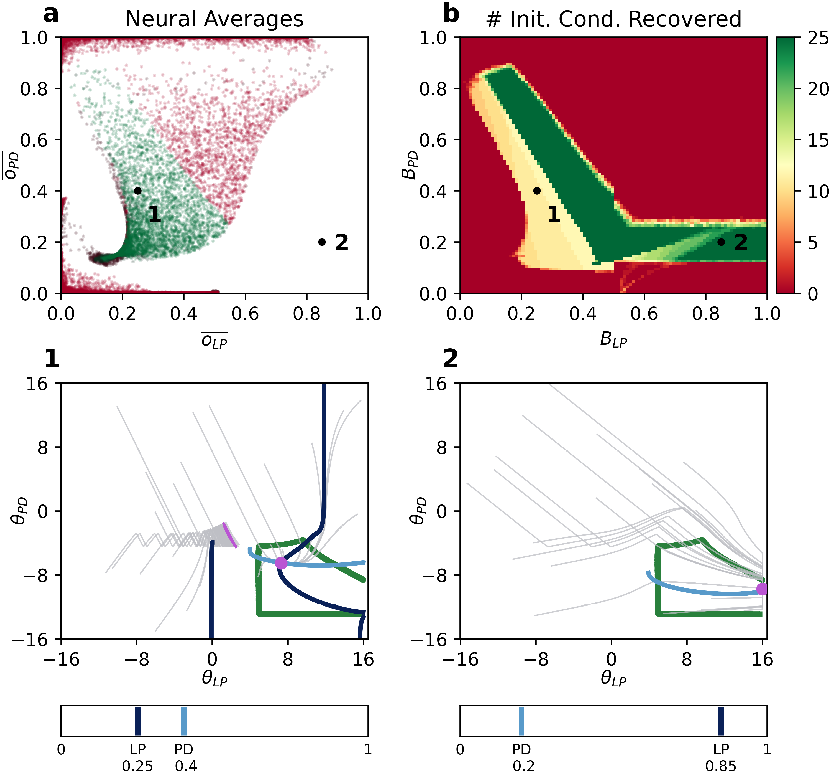
Some circuits represent exceptions to the described predictive framework. The distribution of neural average values in among ADHP-accessible circuit configurations (**a**) is compared to simulated ADHP performance (**b**), as in Fig 6. Two regions of disagreement (points 1 & 2) are attributed to two prominent exceptional phenomena. (**1**)) When ADHP targets are set to match point 1, we predict a single pyloric endpoint. In simulations, randomly initialized trajectories approach either a pyloric endpoint (pink point) or oscillate indefinitely (stable limit cycle outlined in pink). At no point in this limit cycle are target averages stably attained. Instead neural states flip-flop between two fixed points with values above and below their targets, which abruptly appear and disappear as the system is pushed across a pair of bifurcation lines. (**2**)) When ADHP targets are set to point 2, the upper regulatory boundary *θ*_*LP*_ = 16 is exploited to create a pyloric endpoint which attracts all trajectories (pink point), even though 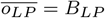 is unattainable.

Other regulators of this circuit demonstrate the occasional importance of ADHP’s interaction with parameter boundaries. In this model, because we restrict neural biases to the range [-16,16], these lines constitute extra nullclines for all ADHP mechanisms, independent of their target activity values. These nullclines are stable if the homeostatic flow vector points outside the boundary, and if they run through the pyloric region, then they can intersect nullclines for other parameters to produce pyloric endpoints. This is the case for our example circuit, which can be robustly regulated by a “stripe” of ADHP mechanisms targeting average PD activity values which exist in the pyloric region at the boundary, and LP activity values greater than those at the boundary (Fig. 7a-b). However these bounds on parameter values arise, whether discretely or asymptotically, emerging from other rules or imposed directly by modelers, they would have the effect of introducing another nullcline. Effective prediction of ADHP performance must therefore consider not only pairs of target averages that exist in the ADHP-accessible plane, but pairs consisting of averages at parameter boundaries and values of the other parameter that exceed its boundary value.

In this section, we have outlined a framework for predicting the success of zero-range ADHP mechanisms in restoring a functional feature of interest (pyloricness) after parametric perturbation of a given circuit. Put simply, ADHP that targets neural activity values which always co-occur with pyloricness will reliably recover pyloricness. Even though average neural activity itself is not a feature of interest, or even a good indicator of that feature in general, it only matters that in relevant subsets of parameter space, average neural activity serves as a reliable *proxy* for functionality. The relevant subset of parameter space, in this context, is the set of circuit configurations reachable by ADHP from the circuit’s perturbed state. So far, these have been 2-dimensional parameter space slices through known pyloric circuits, for which particular ADHP mechanisms may or may not be reliable regulators. Next, we will ask how these insights apply when ADHP must generalize to variety of degenerate circuits occupying different regions of parameter space with different surroundings. Then, we apply our predictive framework to ADHP operating in three dimensions (rather than two) and highlight important considerations for increasing the number of regulated parameters.

### 3.4 Generalization across degenerate circuits

In our study thus far, we have optimized and analyzed the regulatory performance of ADHP for particular, restricted regions of parameter space surrounding optimized pyloric circuits. We have often found that the most successful regulators of one circuit in our database differ from the most successful regulators of another. Presumably, though, ADHP would be most beneficial if it were generalizable across a wide array of circuit configurations, rather than individually tailored. After all, even as organisms develop and differentiate from others of their species, they must be able to rely on some consistent regulatory mechanism to maintain functionality. We will therefore use the present model to ask, under what conditions can particular homeostatic plasticity mechanisms generalize across degenerate circuit configurations?

The most straightforward generalization scenario is when the distribution of average neural activity values in the vicinity of different circuit configurations is similar. Crucially, if the set of neural activity averages that specify pyloricness for one configuration overlaps with the set of neural activity averages that specify pyloricness for another, then a generalized ADHP mechanism can be devised which targets activity averages in this overlap. An example of this is shown in Figure 8a, where the set of zero-range ADHP mechanisms that successfully regulates Circuit 1 overlaps with the set that successfully regulates Circuit 2.

**Fig. 8.**
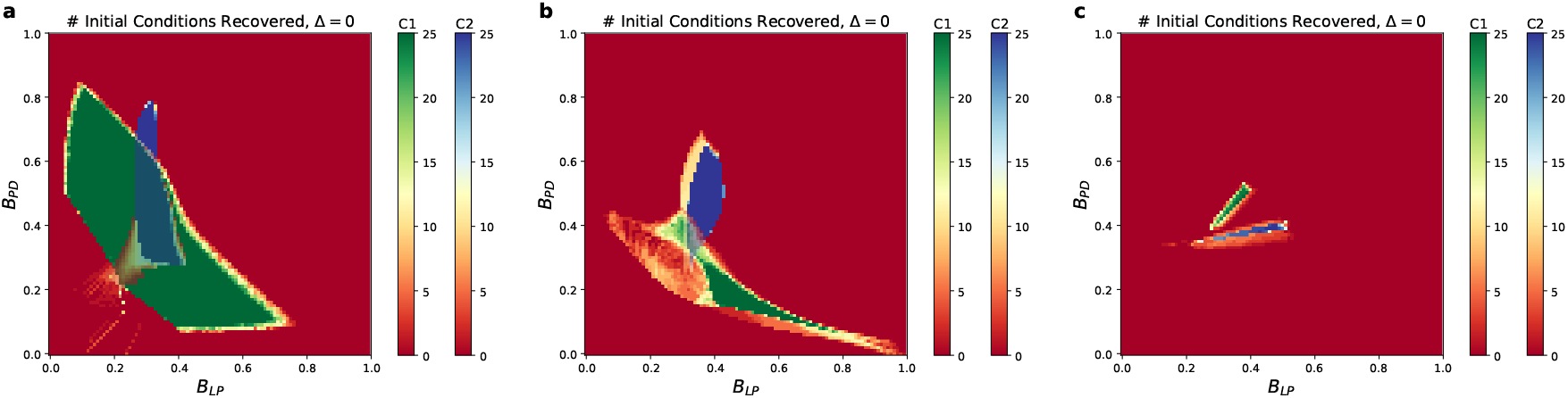
Possibilities for generalization across degenerate circuit pairs. We plot simulated recovery performance throughout ADHP meta-parameter space for pairs of example circuits using two different color scales.**a:** Most pairs (*n* = 3057 of 3160) of circuits in our primary dataset are amenable to generalized regulation, like those pictured. Areas of overlap between solid green and solid blue indicate ADHP mechanisms with those target activity values reliably regulate both circuits. **b:** ADHP can only partially generalize across other circuit pairs. Solid green and solid blue regions have no overlap, but there are some reliable regulators of circuit 2 which can partially regulate circuit 1. **c:** Some circuit pairs defy even partial generalization. No ADHP mechanism which successfully regulates either circuit can generalize to the other.

If candidate circuits do not share any neural averages which uniquely specify pyloricness, our framework suggests that ADHP mechanisms with zero target range cannot reliably regulate them both. Compromises can sometimes be made if there exist ADHP mechanisms which at least partially regulate both circuits (as in Figure 8b). Mechanistically, this means that pairs of neural averages which specify pyloricness in one circuit at least exist within the pyloric region of the other, though they also exist outside it, leading to perturbation-dependent performance. A worse case is when every pair of neural averages specifying pyloricness for one circuit configuration exists only outside the pyloric region that surrounds the other circuit, and vice-versa (Figure 8c). In this case, even though each circuit could be reliably regulated on its own, no ADHP mechanisms can be devised which generalize between them.

All three of these possibilities are realized between pairs of individually regulable circuits in our database of well-timed pyloric oscillators. Surprisingly, a vast majority of circuit pairs are mutually regulatable, with overlapping sets of successful ADHP mechanisms. There are 80 circuits in the database for which at least one reliably successful ADHP mechanism exists, yielding 3160 testable circuit pairs. For 3057 (96.7%) of these pairs, there is overlap in the set of successful regulators, indicating the existence of an ADHP mechanism which could generalize to both circuits. Given the remarkable rate of ADHP generalization in this database of circuits with similar oscillation timing, we wondered how much this result was driven by circuit similarity. We thus repeated the generalization analysis for our other, more heterogeneous, dataset of pyloric circuits. In this dataset, there are 85 circuits which can be reliably regulated by some zero-range ADHP mechanism (3570 testable circuit pairs). Testing for ADHP generalizability across circuit pairs, we found a significantly reduced success rate (2469/3570 = 69.2% of pairs). Thus, it is more likely for a single ADHP mechanism to generalize across circuits with more similar dynamics than those with differing dynamics.

We can more precisely account for this observation using the presented framework, which explains ADHP success using the distribution of pyloric and non-pyloric average activity values in given subset of parameter space. Even though circuits in the well-timed dataset all had different parameters, it seems likely that they come from a relatively limited neighborhood of parameter space. Within this neighborhood, neural average values inside and outside of the pyloric region are more similarly distributed, as compared to their distributions in parameter space as a whole. This would explain why ADHP mechanisms, which operate by specifying pyloric endpoints to the exclusion of non-pyloric circuit configurations, are more likely to generalize among these circuits (see also Fig 11). Our framework also highlights that the question of homeostatic generalizability requires examining the distribution of both functional and *non-functional* circuit configurations in candidate regions of parameter space. The separability of these distributions is what underlies the success of regulatory mechanisms, both for individual circuits and diverse populations.

In this section, we have examined the generalizability of ADHP across circuits which differ in their unregulated parameters (so far, these include neuronal time constants, synaptic weights, and *θ*_*PY*_). And, while it is certainly true that some of the properties of neural circuits are outside the control of any given homeostatic plasticity mechanism, it is unclear how the ratio of regulated to unregulated parameters affects homeostatic prospects. In the next section, we illustrate the consequences of a small increase in this ratio.

### 3.5 Navigating higher-dimensional parameter spaces

As a final extension of this model, we will apply our predictive heuristics to ADHP regulating all three neural biases (*θ*_*i*_, *i* ∈ {*LP, PY, PD}*), expanding the homeostatically accessible subspace from two to three dimensions. The computational burden of calculating the average activity and pyloricness of circuits in a higher-dimensional subspace with the requisite resolution increases exponentially, but even one step in this direction serves to demonstrate essential considerations.

First, given our results in the 2D case, what should we expect about the benefits and drawbacks of bringing another neuron’s bias under homeostatic control? On one hand, it must be beneficial to add another attribute (in this case the average activity of the PY neuron) by which circuit configurations can be distinguished (Olypher & Calabrese, 2007; Onasch & Gjorgjieva, 2020; Yang & Prescott, 2023). Perhaps, for instance, a set of pyloric and non-pyloric configurations which is inseparable on the basis of their average LP and PD activity is separable into pyloric and non-pyloric subsets based on PY activity. Another potential benefit is that a larger ADHP-accessible search space may contain new pyloric circuit configurations to target, some of which may be separable from non-pyloric configurations even if those in the 2D subspace were not. On the other hand, this larger space may also include more *non-pyloric* circuit configurations, with average activity values that spoil previously separable pyloric sets. So, which effect of increased dimensionality is stronger in our population of model circuits?

To answer this, we systematically searched 3D ADHP meta-parameter space (target activity values for each neuron ranging from 0 to 1 in steps of 0.05) for regulators that could maintain pyloricness for each individual circuit in our primary database. We tested each mechanism for its ability to recover pyloricness from 27 different bias perturbations on the 3-by-3-by-3 grid {−10, 0, 10}^3^. We found that the best 3D ADHP almost always met or surpassed the capabilities of 2D ADHP for individual circuits. Each of the circuits whose 2D regulation was limited by the inseparability of pyloric and non-pyloric averages could be perfectly regulated by a 3D ADHP mechanism, suggesting separability of functional and nonfunctional configurations along the third dimension. Even the circuit which completely defied maintenance by 2D ADHP could be perfectly regulated by 3D ADHP. Indeed, there was only one circuit out of 100 for which no 3D ADHP mechanism could perfectly recover pyloricness. Generally, it appears the benefits of increasing homeostatic dimensionality for these circuits outweigh any drawbacks.

It was unclear, however, whether the same necessary and sufficient conditions that proved predictively valuable in 2D were still valid in higher dimensions. To assess this, we extended the prior methods to predict the performance of 3D ADHP mechanisms on one of our evolved circuits, and compared these to simulation results. The circuit being regulated in this analysis is the second example circuit from figures 2-6 above. 2D ADHP mechanisms could not robustly regulate this circuit because pyloric circuit configurations in the 2D subspace were inseparable from non-pyloric ones. We simulated 1,000,000 3D ADHP mechanisms with target activity levels of all three neurons on an evenly-spaced 3D grid, ranging between 0 and 1 in steps of 0.01. As before, the performance for each was assessed by the proportion of parametric perturbations from which the circuit was recovered to pyloricness after 50,000 seconds simulation time (Figure 9a). 3D ADHP mechanisms across a substantial range of target activity values were found to have perfect performance.

**Fig. 9.**
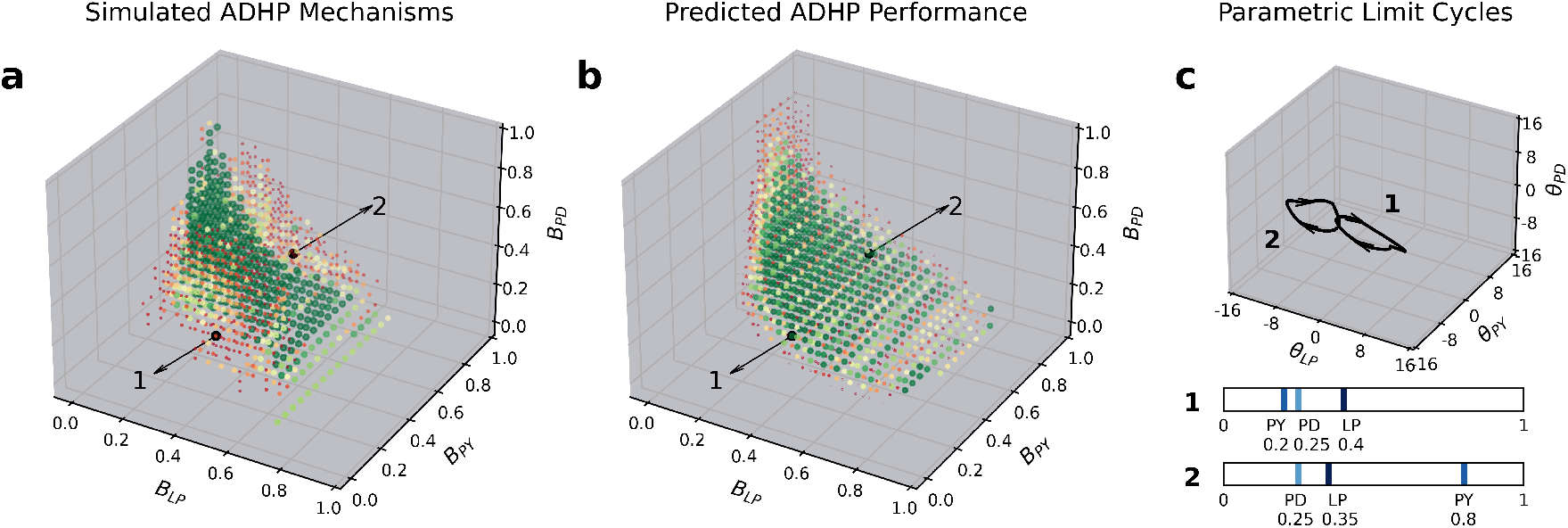
Our predictive framework for ADHP performance can be extended to three regulated dimensions (*θ*_*LP*_, *θ*_*PY*_, and *θ*_*PD*_). **(a)** We systematically simulate 1,000,000 ADHP mechanisms, with the target activity levels of all three neurons defined on a 3D grid, ranging between 0 and 1 in steps of 0.01. Color (red to green) and size (small to large) of shown points reflects the nonzero number of parametric perturbations (of 27 selected on the grid {−10, 0, 10}^3^) successfully recovered to pyloricness by the corresponding ADHP mechanism after 50,000 seconds (13.9 hrs) of simulated time. **(b)** We compare this to the predicted performance for each ADHP mechanism. Here point color and size reflects the ratio of pyloric to nonpyloric ADHP-accessible circuit configurations (as sampled on the 3D grid where each neural bias ranges from −16 to 16 in steps of 0.05) with average neural activity values that most closely match those targets. Large green dots mark those average values which only occur in the pyloric region of the 3D ADHP-accessible subspace. Small red dots mark those which occur most always in the non-pyloric region, and those which never occur in the pyloric region are not pictured. **(c)** We highlight two representative ADHP mechanisms, 1 and 2, from regions of meta-parameter space where simulated performance falls short of predictions. We see that parameter space trajectories converge to the limit cycles labeled 1 and 2, which have periods of about 8575 and 8150 seconds, respectively.

**Fig. 10.**
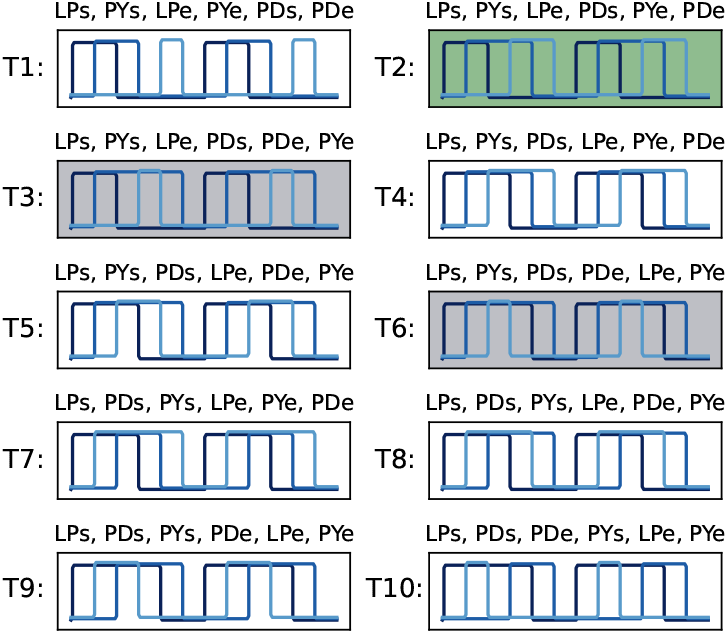
Pyloric Oscillation Types. Among rhythms where each neuron (LP, PY, PD) has one defined burst per cycle, there are 5! qualitatively different ways to order the beginning and end of each burst. Ten of these orderings meet all pyloric criteria set out in Section 2.1, with standardized schematics pictured here. Model circuits within the same category may exhibit varied duty cycles and different values during bursting/up and quiescent/down epochs.

**Fig. 11.**
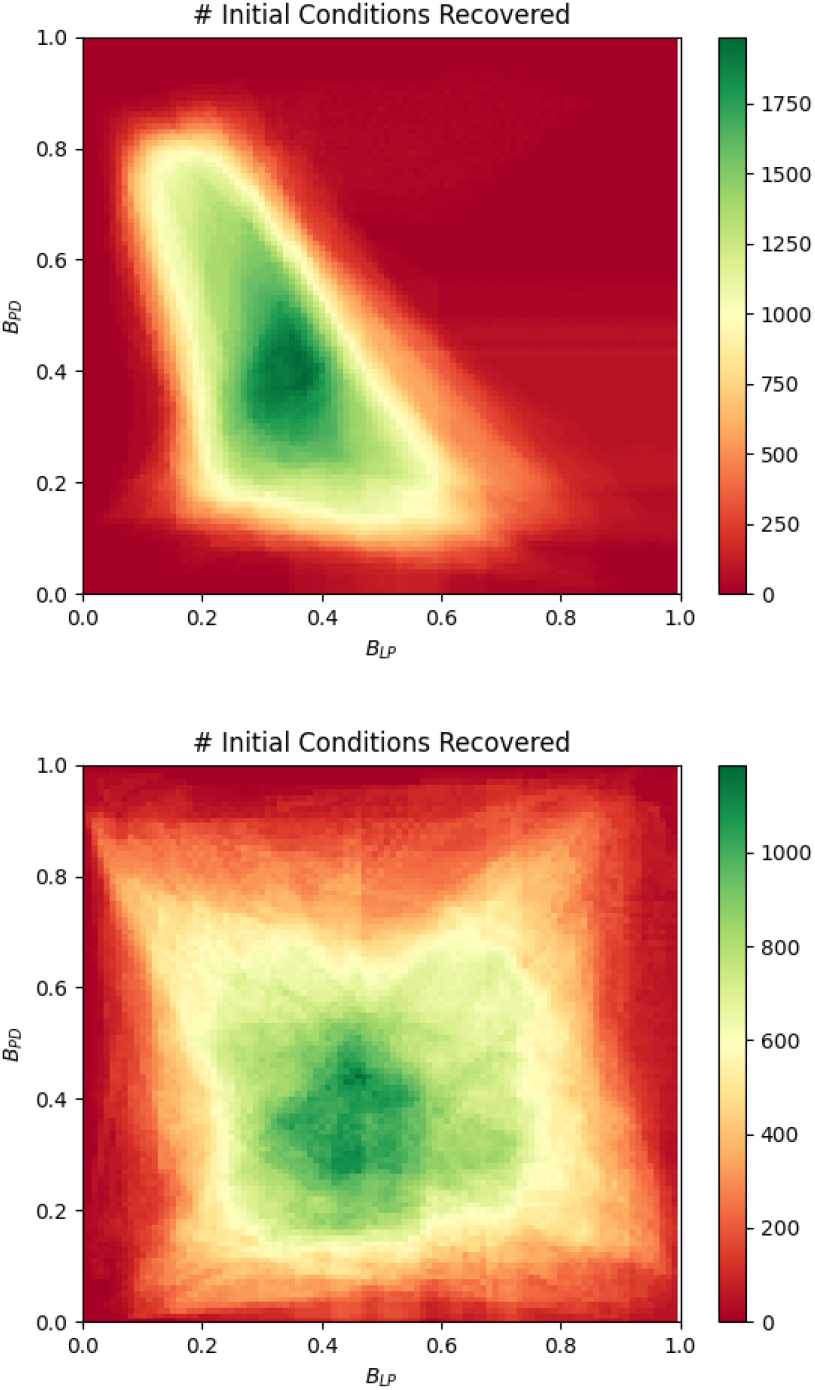
Aggregate of recovery performance for all zero-range ADHP mechanisms, across all evolved pyloric circuits. Mechanisms are tested for their ability to recover pyloricness within 50,000 seconds following 25 perturbations to each of 100 circuits in both datasets (max performance = 2500). **Top:** Results on the more homogeneous evolved dataset are highly consistent, producing a recognizable shape that mirrors ADHP performance on many individual circuits (e.g. Fig 6b). **Bottom:** Results from the more heterogeneous dataset (evolved without timing rewards) are relatively inconsistent, but successful mechanisms tend to have low-moderate activity targets for both neurons, as we would expect for neurons in a tri-phasic oscillation.

We then checked whether observed performance occurs where we expect it to, based on the distribution of average neural activity values among pyloric and non-pyloric circuits in the 3D accessible subspace. As before, we sample circuit configurations throughout this subspace, holding neural time constants and connection weights at evolved values but varying neural bias values on an evenly spaced 3D grid, ranging from −16 to 16 in steps of 0.05. We let neural states equilibrate for each circuit configuration before evaluating its pyloric status and the time-averaged activity values for each neuron. For visual clarity and accordance with simulation procedures, we round the average activity values to the nearest 0.01 and group each circuit configuration with others of the same rounded averages. Then, for each set of rounded activity values, we can evaluate what percentage of circuit configurations exhibiting those averages are pyloric and non-pyloric. The result is illustrated in Figure 9b. We find there is a strong accord between predicted and simulated pyloric recovery by 3D ADHP mechanisms on this particular circuit. There is indeed a central concave volume in ADHP meta-parameter space which captures high-performing mechanisms with activity targets that are uniquely (or nearly uniquely) represented among pyloric circuit configurations.

There are, however, two regions of ADHP meta-parameter space where simulated performance falls short of predictions. To account for this discrepancy, we take a closer look at particular ADHP mechanisms in these regions. We see that the corresponding homeostatic trajectories do not approach single points in neural parameter space, nor oscillate around them with small amplitude. They maintain stable, large-amplitude oscillations which often pass through pyloric and non-pyloric regions (see Fig 9c). When this is the case, as it is for these two example mechanisms, the pyloric character of the neural state dynamics depends on the phase of the oscillation. Even though our framework predicts suggests that all homeostatic attractors in the ADHP-accessible subspace are within the pyloric region, homeostatic trajectories instead follow this attractive cycle. Long-timescale limit cycles also explain the newfound regulatability of the previously unstable circuit (third example circuit in Figures 2-6). We know that all the pyloric circuit configurations in the 2D subspace around this circuit are structurally unstable. And it is not the case that the 3D subspace contains stable configurations which can be reliably targeted by ADHP. Instead, 3D ADHP mechanisms are capable of inducing a long-timescale limit cycle which is fully contained in the pyloric region. Even though parameters oscillate indefinitely, neural dynamics remain pyloric.

In fact, long-timescale limit cycles like these cannot exist unless there is some region of the ADHP-accessible parameter subspace which exhibits homeostatic instability in at least one dimension. That is, there must be at least one regulated parameter which, for some circuit configurations, has the opposite of its “intended” effect on average neural activity. This situation, of course, is increasingly likely with the addition of more regulated dimensions.

Overall, when these limit cycles are present, they often affect ADHP performance predictions. In their absence, ADHP performance is still well predicted by the distribution of neural activity values among pyloric and non-pyloric circuits in the ADHP-accessible subspace. Future studies might draw from bifurcation theory to pinpoint specific properties of ADHP mechanisms (and the landscapes in which they operate) which transform homeostatic endpoints into parametric limit cycles. Combined with the current framework, this would provide a complete account of ADHP outcomes which scales well many regulated dimensions.

## 4 Discussion

Using our model of the pyloric central pattern generator, we have demonstrated the conditions required for local activity-dependent homeostatic plasticity to regulate circuit-level dynamic properties. We found that regulatory success depends on a nexus of features involving the initial circuit configuration, homeostatic neural activity targets, and perturbations to be recovered. Recovery is most reliable when, within the set of circuit configurations accessible from the perturbed state, satisfaction of homeostatic targets always implies higher-level functionality, while all nonfunctional circuit configurations produce homeostatic change. Under these conditions, barring other factors like boundary interactions, long-timescale oscillations, and attractive bifurcation points, any perturbation of regulated parameters will be recovered to a functional state. Otherwise, non-functional configurations which satisfy homeostatic targets become unwanted attractors, making recovery outcomes perturbation-dependent.

The maximum ability of ADHP to regulate a circuit varies based on the circuit’s configuration. Some configurations, despite producing valid pyloric oscillations, cannot be reliably maintained by any ADHP mechanism. We have highlighted several potential causes of this in our pyloric model. For one, it may be that, within the parameter subspace defined by an initial circuit and all possible perturbations of the regulated parameters, there is no set of neural activity values which specifies functionality (pyloricness) while excluding all non-functional activity (non-pyloricness). Pyloric activity averages, in other words, are *inseparable* from non-pyloric ones. Therefore, even in the best case scenario, homeostatic recovery would be perturbation-dependent. Alternatively, any distinguishable pyloric configurations which do exist may be unstable with respect to regulatory rules. While it makes intuitive sense that ADHP should increase individual neural excitability levels to bolster diminished activity (and vice versa), we have shown that for certain circuits this may actually have the opposite effect. Homeostatic trajectories will be repelled from all pyloric circuit configurations for which this is true. These conditions highlight the fact that, to adequately explain homeostatic performance, we must understand both the operation of homeostatic mechanisms and the parametric landscape on which they operate.

Having described this quality of being relatively “easy” or “hard” to regulate, we may also consider it a selection criterion in itself. In order to cultivate robust neural circuits, evolution and development may select for configurations which are easy to maintain with available regulatory mechanisms. In the context of the current model, this means favoring circuit parameters where pyloric and non-pyloric configurations in the ADHP-accessible subspace have clearly distinguishable neural averages. In subspaces where this is not the case, it is impossible to tune any ADHP mechanism which reliably avoids dysfunctional states. Thus, even though such parameter sets could produce perfectly valid pyloric oscillations, they may not ever be observed in real animals because they are difficult to maintain. We would predict, moreover, that those configurations which *are* observed share certain characteristics that correlate with regulatability.

Our model makes several other experimentally testable predictions. For example, we have seen ADHP often benefits from having specific and restrictive activity targets, thereby destabilizing the greatest number of dysfunctional configurations. If this is the case, then we expect real ADHP mechanisms to be able to discriminate between (differentially respond to) relatively similar activity levels when sustained for sufficient time. Other hypotheses about homeostatic targets could be adjudicated with time-varying data on neural conductances. If neural conductance values oscillate with small amplitude in phase with rhythmic neural activity, then ADHP’s target activity range is likely centered within the range of values those neurons take on. Conversely, if neural parameters oscillate slowly with large amplitude, then none of the activity values the neurons take on during the cycle corresponds exactly to homeostatic targets. Finally, we emphasize that, among the wide selection of circuit configurations with desirable dynamics, ADHP need only stabilize a subset. This means that perfectly functional configurations may be passed over during the process of recovery. Indeed, this appears to occur during observed “bouts”, where recovering pyloric circuits alternate between epochs of pyloric activity and non-pyloric activity before eventually stabilizing (Luther et al., 2003; Zhang & Golowasch, 2011). If it is true that bouts indicate brief forays through the pyloric region in the recovering circuit’s parametric trajectory, then the activity patterns observed during these bouts should differ in principled ways from the activity observed at steady state. In particular, they should differ with respect to ADHP control variables (i.e. time-averaged activity). Investigation of these hypotheses would greatly elucidate the way that underlying parameter space structure, perturbations, and homeostatic mechanisms interact to produce observed variability in functional neural circuits.

We also suggest some worthwhile extensions of this particular model which could help bridge the gap to experimental data. For instance, given the apparent importance of regulatory specificity in the exclusion of non-pyloric steady states, one could examine the effects of noise on homeostatic outcomes. Sources of noise, either in neural states or ADHP’s detection of them, may decrease discriminatory capacity and negatively impact function. On the other hand, since only the average values of time-varying neural states produce sustained homeostatic activation, noise centered around zero might *not* appreciably affect ADHP. Also, we note that the heuristics derived for predicting the recoverability of perturbations rely on the fact that ADHP can tune parameters independently of one another. Each model neuron’s activity was used to tune exactly one parameter (*θ*_*i*_), and therefore movement in any direction through the ADHP-accessible parameter subspace was allowed. In real neural circuits, however, ADHP simultaneously regulates multiple parameters associated with each neuron, and some of these processes even share molecular machinery (MacLean et al., 2003, 2005; Northcutt & Schulz, 2019; Pozo & Goda, 2010; Ransdell et al., 2012; Temporal et al., 2014). As such, homeostatically induced changes for multiple parameters are tied together, and movement through parameter space is restricted to mechanistically defined directions. Other ADHP models which incorporate this interdependence demonstrate that it limits the set of ADHP-accessible circuit configurations from any given state, making regulatory outcomes more perturbation-dependent (Abbott & LeMasson, 1993; Franci et al., 2020; LeMasson et al., 1993; Olypher & Prinz, 2010), and possibly imposing correlation structure among steady state solutions (O’Leary et al., 2013). Future iterations of this model could traverse the spectrum of increasingly interdependent parameter regulation and incorporate its effects into the present predictive framework.

Moreover, our methods are not specific to this instantiation of ADHP or the pyloric pattern generator. Many of our observations and analyses could be extended to other models so long as certain conditions are met. ADHP, for example, could be implemented with alternative activity detection schemes, with different control architectures, or even as a mechanistically grounded rate-based model. It matters only that it can change system parameters, and that the sign of this change can be predicted by measuring some control variable which is defined for all circuits. Likewise, the circuit’s function or property of interest can be defined in any way, and may indeed be quite abstracted from individual neural activity levels. Instead of pyloricness, one might consider a circuit’s ability to perform computations over some input range, interface with other parts of the nervous system, detect stimuli, or even a combination of such features. So long as the function of interest differs from the ADHP control variable and is well-defined for all circuit configurations, one can delineate the set of reachable circuit configurations from any perturbed state, divide it into subsets of functional and non-functional configurations, and predict regulatory outcomes by analyzing the overlap between values of the homeostatic control variable observed in each subset.

In sum, our model has generated valuable insights about the ability of local ADHP to maintain circuit-level properties. The most important condition for robust homeostasis is that, within the set of neural activity patterns expressed by ADHP-accessible circuits, only desirable ones are balanced over ADHP’s target range, while undesirable patterns induce net parameter change. Even if this is impossible to guarantee when considering all available circuit configurations, it will be possible within certain subsets of parameter space. Well-tuned ADHP operating on circuits within these subsets can reliably rescue function after parametric perturbations. In other subsets of parameter space where it is impossible to cleanly distinguish functional and dysfunctional neural activity from local information alone, ADHP’s success cannot be guaranteed. Simply by examining the landscape of possible functional and dysfunctional circuit configurations, we can generate hypotheses about homeostatic mechanisms, and predict recovery outcomes across individuals.

## Appendix

## Declarations

### Conflict of Interest Statement

The authors have no competing interests to declare.

### Code Availability

C++ and Python code required to reproduce all data can be found at https://doi.org/10.5281/zenodo.18509900

## Acknowledgments

We would like to thank Connor McShaffrey, Denizhan Pak, and Eden Forbes for their feedback on this manuscript. This research was supported Lilly Endowment, Inc., through its support for the Indiana University Pervasive Technology Institute.

## Notes

https://doi.org/10.5281/zenodo.18509900

### Competing Interest Statement

The authors have declared no competing interest.

